# Bacterial Vipp1 and PspA are members of the ancient ESCRT-III membrane-remodelling superfamily

**DOI:** 10.1101/2020.08.13.249979

**Authors:** Jiwei Liu, Matteo Tassinari, Diorge P Souza, Souvik Naskar, Jeffrey K. Noel, Olga Bohuszewicz, Martin Buck, Tom A. Williams, Buzz Baum, Harry H Low

## Abstract

Membrane remodelling and repair are essential for all cells. Proteins that perform these functions include Vipp1/IM30 in photosynthetic plastids, PspA in bacteria, CdvB in TACK archaea and ESCRT-III in eukaryotes. Here, we show that these protein families are homologous and share a common evolutionary origin. Using cryo-electron microscopy we present structures for Vipp1 rings over a range of symmetries. Each ring is built from rungs that stack and spontaneously self-organise to form domes. Rungs are assembled from a polymer that is strikingly similar in structure to ESCRT-III. A tilt between rungs generates the dome-shaped curvature with constricted open ends and an inner membrane-binding lumen. Overall, our results reveal conserved mechanistic principles that underlie Vipp1, PspA and ESCRT-III dependent membrane remodelling across all domains of life.

**One sentence summary:** Evolutionary and structural analyses of Vipp1/IM30 rings reveal ESCRT-III-like polymers that remodel membranes in bacteria.

## Main text

Various cytoskeletal elements, filaments and membrane remodelling systems that were once viewed as the defining features of eukaryotic cells are now known to have a prokaryotic ancestry. Bacteria and some archaea have their own versions of tubulin (FtsZ), actin (MreB, FtsA and ParM), intermediate filaments (Crescentin), and dynamins (BDLPs) (*1*-*3*). These proteins perform functions in prokaryotes that are often analogous to their eukaryotic counterparts including cell division, cell shape control and membrane remodelling. In contrast, homologues of ESCRT-III proteins, another conserved and important superfamily of membrane remodelling proteins in eukaryotes, which perform diverse functions in multivesicular body formation, cytokinetic abscission, plasma membrane repair, nuclear envelope reformation and viral budding (*4*), have yet to be found in bacteria. The recent identification of ESCRT-III homologues in TACK (*5, 6*) and Asgard (*7*) archaea where they have been proposed to perform important roles in viral budding, exosome formation and cell division suggested that these eukaryotic signature proteins likely have an archaeal origin.

### PspA, Vipp1 and ESCRT-III are close evolutionary relatives

To identify other as yet unknown ESCRT-III relatives, we used sensitive protein sequence searches based on Hidden Markov Models (*8*). Using eukaryotic ESCRT-III proteins as search queries, PspA and Vipp1/IM30 families were identified as bacterial ESCRT-III homologues (materials and methods and fig. S1A). This was intriguing as PspA functions in the bacterial membrane stress response and in membrane repair (*9*), whilst Vipp1 in cyanobacteria and chloroplasts functions in thylakoid biogenesis and membrane repair (*10*) through poorly understood mechanisms. These roles are consistent with some of the reported ESCRT-III functions in eukaryotes (*11*). Both PspA and Vipp1 bind membrane and self-assemble to form rings and cage-like scaffolds (*12, 13*). How polymerisation is regulated and coupled to membrane repair in these systems remains a key question.

In addition to their similarity in primary sequence, PspA, Vipp1 and ESCRT-III families were also found to share similar secondary structure predictions with each containing six helices, termed helix *α*0-*α*5. An additional C-terminal extension that includes helix *α*6 is also often present in members of PspA/Vipp1 and ESCRT-III family proteins (Fig. 1A and fig. S1A). Using a co-evolutionary analysis to identify tertiary structural features of the proteins (*14, 15*), we identified high scoring residues that co-vary in PspA/Vipp1 evolution within a key conserved ESCRT-III interface that forms between helix *α*5 and helices *α*1 and *α*2 (Fig. 1B). This interaction functions to maintain ESCRT-III in its auto-inhibited closed monomeric conformation (*16*) and undergoes a helix *α*5 domain swap between subunits to switch the ESCRT-III monomer to an open conformation during polymer assembly (*17*). Overall, a co-evolutionary model of Vipp1 structure based on residue-residue distance constraints was congruent with the known structure of ESCRT-III CHMP3 (Fig. 1B). An analysis mapping the presence or absence of these proteins within bacteria and archaea revealed that PspA/Vipp1 is distributed across bacteria (Fig. 1C) - where ESCRT-III homologues appear completely absent. Conversely, ESCRT-III homologues clustered amongst the archaea within Asgard and TACK superphyla (*7*) (Fig. 1C). In addition, PspA/Vipp1 homologues were found within the euryarchaeata (including in Haloarchaea and Methanosarcinia) where they appeared to have been acquired by horizontal gene transfer from bacteria (Fig. 1C and 1D). A broader phylogenetic analysis of the PspA, Vipp1 and ESCRT-III superfamily as a whole revealed a long branch separating ESCRT-III and PspA/Vipp1 clades, indicating a divergence between these two subfamilies early in the evolutionary history of life (Fig. 1D and fig. S1B). As the long branch at the centre of the phylogeny corresponds to the basal divergence between archaea and bacteria, and the archaeal *pspA* genes were likely acquired from bacteria, the tree is consistent with the hypothesis that the divergence between PspA and ESCRT-III occurred in the last universal common ancestor (LUCA). This analysis also confirmed that Vipp1 likely arose from a *pspA* gene duplication (*18*) and is found widely distributed amongst the cyanobacteria and plants, often with multiple copies. Some cyanobacteria also have PspA with a likely function in cell membrane repair rather than thylakoid membrane maintenance as for Vipp1 (*10*). In order to further develop our evolutionary analyses, and to probe PspA/Vipp1 function and mechanism, structural studies were undertaken focussed on Vipp1.

**Fig. 1.**
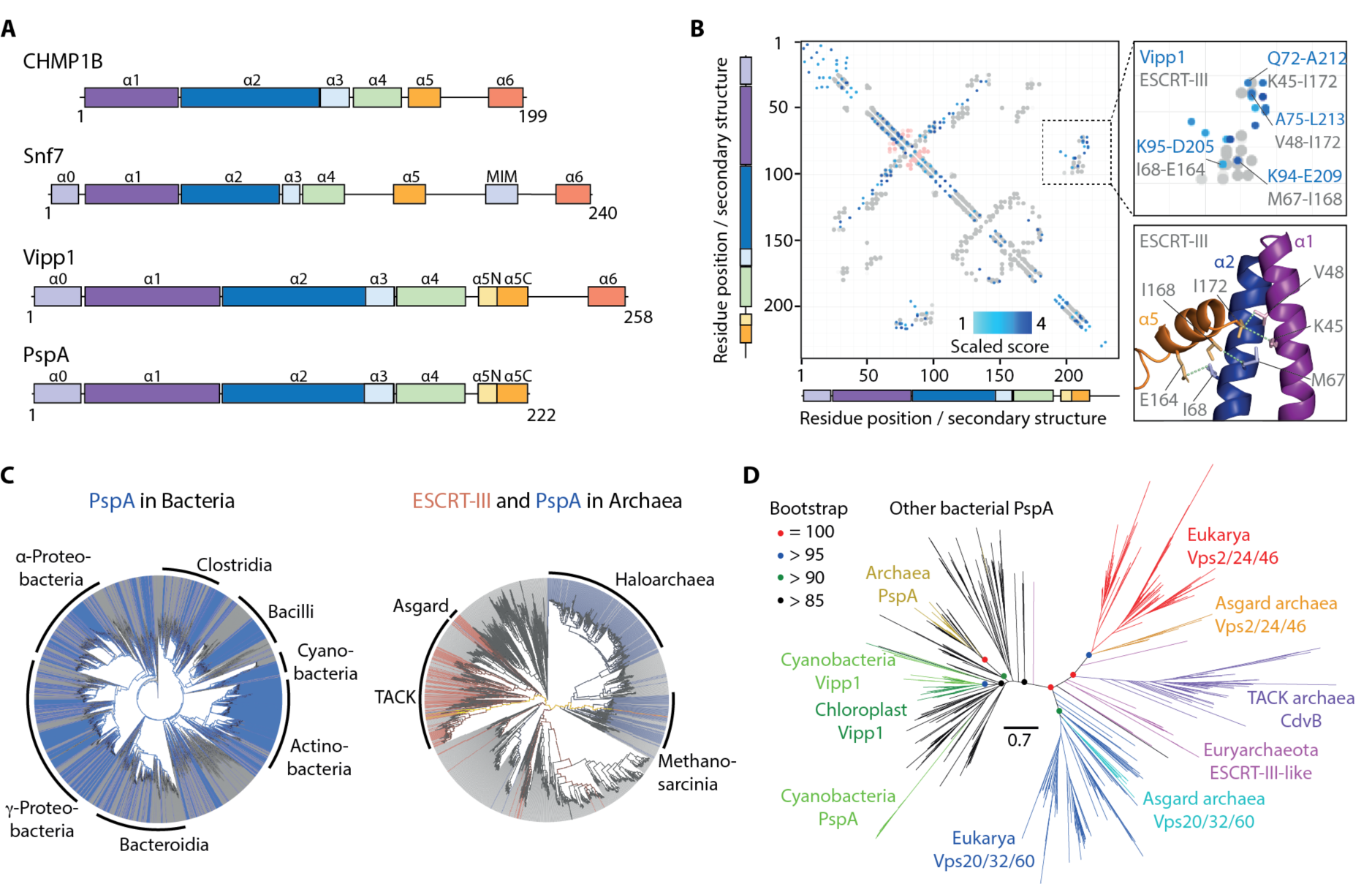
ESCRT-III/Snf7/CdvB and PspA/Vipp1 proteins are part of the same super-family. (**A**) ESCRT-III (human CHMP1B and yeast Snf7) and PspA (*N. punctiforme* Vipp1 and *E. coli* PspA) protein families are homologues (Fig S1A) and have a similar secondary structural organisation (alpha helices labelled *α*0 to *α*6). (**B**) A co-evolutionary analysis reveals similarities in the tertiary structure of PspA and ESCRT-III proteins. (Left) Plot shows a co-evolutionary residue contact map (using numbering based on the *N. punctiforme* Vipp1 sequence) super-imposed on ESCRT-III residue-residue distance data extracted from the experimentally determined structure of human CHMP3 (PDB code 3FRT). Extent of blue shading represents the strength in covariance between Vipp1 co-evolving residue pairs. Grey and red circles indicate intramolecular and intermolecular contacts (< 5 Å) in the CHMP3 X-ray crystal structure, respectively. (Top right) Inset focuses on selected Vipp1 evolutionary coupled residues in blue clustering with known contacts derived from CHMP3 in grey. (Bottom right) These contacts are mapped onto CHMP3 structure. (**C**) A phylogenetic analysis reveals a broad distribution of PspA/Vipp1 (blue) and ESCRT-III (red) homologues across bacteria (left; over 27000 genomes) and archaea (right; over 1500 genomes). Only a very few genomes encode both proteins (yellow). Genomes lacking both PspA/Vipp1 and ESCRT-III are presented in grey. (**D**) Tree of the PspA/ESCRT-III superfamily coloured according to phylogenetic taxons. A long branch separates the PspA/Vipp1 (left) and ESCRT-III (right) subfamilies. Scale bar represents expected substitutions per site.

### Vipp1 purification and cryo-electron microscopy

Full-length Vipp1/IM30 (amino acids 1-258, 28.7 kDa) from *Nostoc punctiforme* was expressed in *Escherichia coli* as a C-terminal maltose-binding protein (MBP) fusion. MBP-Vipp1 was purified by amylose affinity chromatography and then separated from MBP by TEV cleavage and size exclusion chromatography. The purified protein was analysed by SDS-PAGE (fig. S2A) and its identity confirmed by LC-MS analysis. Negative stain (NS) electron microscopy (EM) analysis of Vipp1 revealed a remarkable array of polymeric assemblies, including rings, ring stacks, filaments, and ribbons reminiscent of Vipp1 in other systems (fig. S2B) (*12, 19*-*21*). In order to determine the architecture of the Vipp1 rings, the sample was flash frozen on holey carbon grids for cryo-electron microscopy analysis. Micrographs of Vipp1 in thin vitreous ice revealed preferential orientation bias with an absence of essential side views. In order to promote side view orientation, we mixed pre-formed rat Dynamin 1 filaments with Vipp1 before vitrification. These filaments have a 37 nm diameter and appear equivalent to the human Dynamin 1 super-constricted state (*22*). The resultant network of Dynamin 1 filaments on the grid helped maintain a sufficiently thick ice layer around the Vipp1 particles so that orientation bias was reduced and side views captured (fig. S2C-D). 2D class averages revealed a spectrum of seven ring symmetries ranging from C11 to C17 with symmetries C13, C14 and C15 the most populous (fig. S2E).

### The Vipp1 monomer has an ESCRT-III-like fold

Reconstruction of the C14 symmetry ring (Vipp1_C14_) achieved an overall resolution of 6.5 Å with highest resolution regions reaching 4.8 Å in the central rungs 3-5 (fig. S3 and S4A, Table S1). This resolution was sufficient to allow unambiguous main chain model building of the Vipp1 monomer and asymmetric unit, and consequently the entire 2.4 MDa Vipp1_C14_ ring containing 84 subunits (Fig. 2, materials and methods and fig. S4B-D). To build the Vipp1_C14_ monomer, a homology model was initially generated from the PspA crystal structure hairpin motif (aa 24-142). For these amino acids, PspA and Vipp1 share 32.5 % sequence identity and 58 % similarity with zero gaps, which indicates a conserved sequence register in this section (fig. S4E and fig. S5). The Vipp1 hairpin homology model with its distinct axial twist closely fitted the Vipp1_C14_ map and acted as an anchor from which to build the remaining main chain. Reconstructions were also generated for the other six symmetries and complete ring models built. Sizes ranged from the 1.6 MDa Vipp1_C11_ ring with 55 subunits to the 3.4 MDa Vipp1_C17_ ring with 119 subunits.

**Fig. 2.**
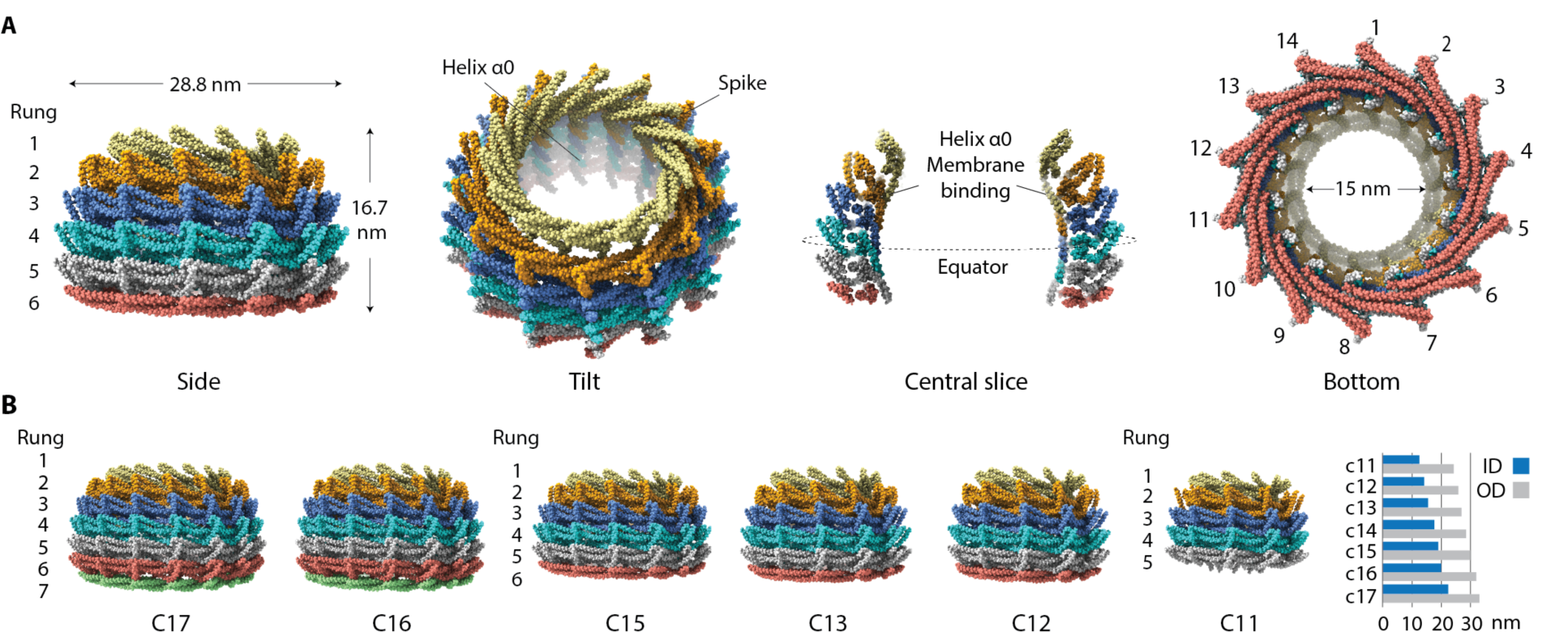
Main chain structures of Vipp1_C11-C17_ rings. (**A**) Structure of Vipp1_C14_. The ring comprises a stack of six rungs. Domed shaped curvature is observed from the side and in the central slice. The ring is widest at the equatorial plane both internally and externally. (**B**) Side view gallery of Vipp1_C11-C17_ rings. Bar chart indicates maximal outer diameter (OD) and maximal inner diameter (ID) as measured at the external and internal equator. Note that Vipp1_C16-C17_ have seven rungs whilst Vipp1_C11_ has just five.

In order to validate these Vipp1_C11-C17_ structures, the accuracy of the sequence register within the Vipp1 monomer was tested by a cysteine cross-linking assay (fig. S6A). L86C and L193C were chosen for mutation (Vipp1_L86C/L193C_) as they are located at opposing ends of the Vipp1 subunit within the hairpin tip and helix *α*5, respectively. Our Vipp1_C11-C17_ structures position these residues at an inter-subunit contact with the cysteine sulphur atoms predicted ∼6 Å apart. In the presence of the oxidising agent ortho-Cu(II)-phenanthroline (CuP) or the cross-linker MTS4 with a 7.8 Å span, clear band shifts were observed for Vipp1_L86C/L193C_ consistent with Vipp1 dimers, trimers, tetramers and higher molecular weight polymers. The subsequent addition of DTT abolished these band shifts due to reduction of the reversible disulphide bond. No equivalent band shifts were observed for the single cysteine mutants Vipp1_L86C_ or Vipp1_L193C_. Overall, this indicates that the Vipp1_L86C/L193C_ band shifts represented specific close cross-links connecting L86C and L193C as predicted by the Vipp1_C11-C17_ structures.

In order to compare the structure of Vipp1 with the structure of the ESCRT-III protein CHMP1B, we followed the ESCRT-III nomenclature (fig. S5) (*17*). The hairpin motif, which is formed by helix *α*1 and conjoined helices *α*2/*α*3, was observed to be a conserved hallmark of Vipp1, PspA and ESCRT-III proteins (Fig. 3A). In addition, helix *α*4 was separated from hairpin helices *α*2/*α*3 by a linker region that corresponded to the ‘elbow’ in CHMP1B (*17*). Helix *α*5 was then angled at near right angles from helix *α*4 in both Vipp1 and ESCRT-III proteins. In addition to these core common helices *α*1-*α*5 between Vipp1 and CHMP1B (*17, 23*), the Vipp1 monomer includes an N-terminal helix *α*0 (aa 1-22) that extends perpendicular to the hairpin and mediates membrane binding in both PspA and Vipp1 systems (*24*-*27*). Helix *α*0 is not observed in CHMP1B, but it is shared by ESCRT-III proteins such as yeast Snf7 where it is reported to also mediate membrane binding (Fig. 1A and fig. S5) (*28*). Vipp1 is distinguished from PspA by an additional C-terminal extension (aa 220-258) comprising a flexible linker and predicted helix *α*6 (*29*). While the C-terminal extension has been shown to negatively regulate Vipp1 self-association *in vivo* (*29*) and might constitute a second lipid binding domain capable of modulating membrane fusion (*21, 30*), it did not yield a density in the Vipp1_C14_ map, suggesting that this region is highly flexible (Fig3A and fig. S4). Interestingly, in the context of ESCRT-III proteins, similar C-terminal extensions that include *α*-helical elements have been shown to function as protein-protein interaction motifs (MIM) (*4*). For CHMP1B, its C-terminal extension binds the ESCRT-III homologue Ist1 to form a dual stranded copolymer (*23*).

**Fig. 3.**
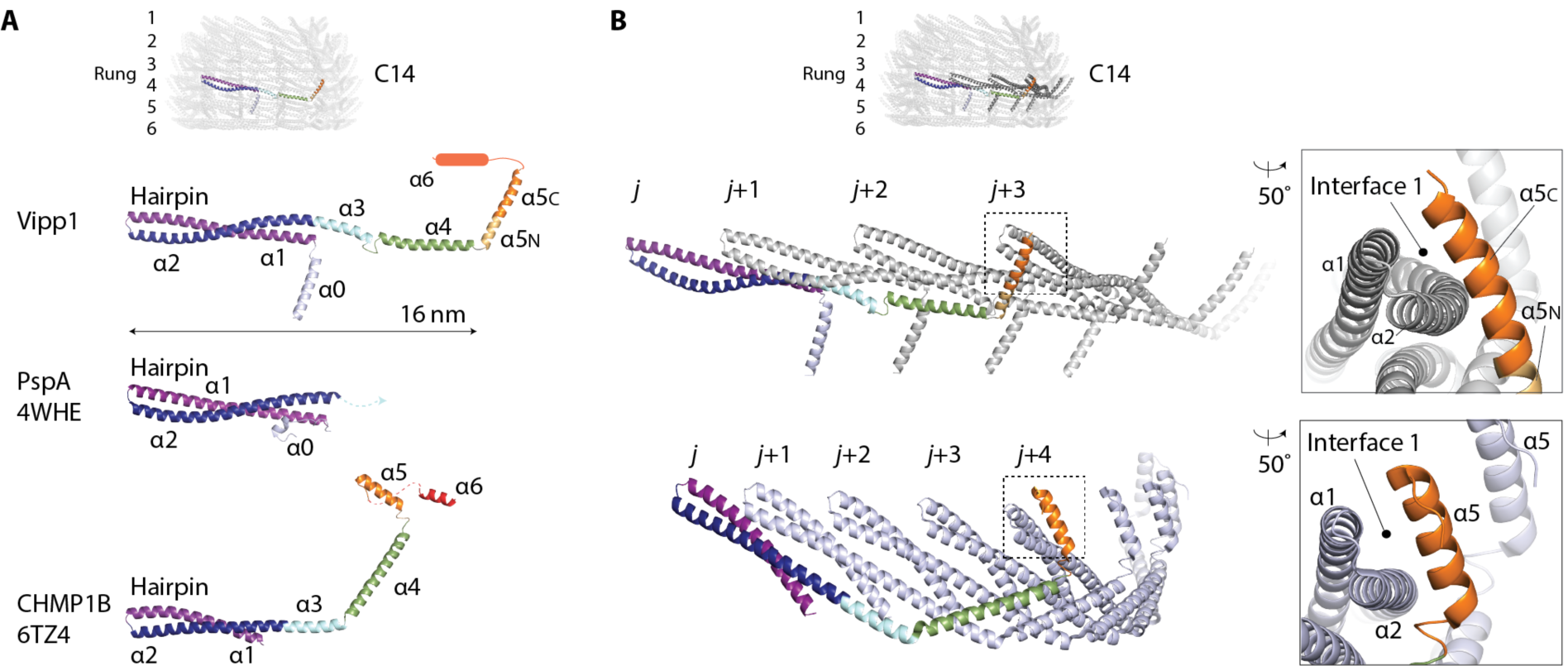
Vipp1 has a similar fold and assembly mechanism as ESCRT-III. (**A**) Vipp1 has the same helical domain organisation as ESCRT-III CHMP1B. The hairpin motif is a hallmark of the Vipp1, PspA and ESCRT-III families. (**B**) Vipp1 and CHMP1B form similar polymers based on hairpin packing and helix *α*5 domain swap. A single circular polymer forms each rung of the Vipp1 ring. Zoom boxes show Interface 1 where the helix *α*5 domain swap binds the hairpin tip of a neighbouring subunit in both Vipp1 (*j*+3) and CHMP1B (*j*+4).

### Vipp1 rungs form ESCRT-III-like filaments

Vipp1_C14_ rings are assembled from six rungs stacked on top of each other (Fig. 2A). Remarkably, each rung appears to regulate the conformation of its neighbour so that a unique asymmetric dome-shaped curvature is achieved when viewed from the side. Internally, rungs are maximally constricted at the top (14 nm) and bottom (15.7 nm) and widest at the dome equator between rungs 3 and 4 (17.5 nm) (Fig. 2A and 2B). This unique asymmetric curvature was a feature of all Vipp1_C11-C17_ ring symmetries. Whilst Vipp1_C12-C15_ rings comprised six rungs, the smallest diameter ring Vipp1_C11_ incorporate five rungs and the larger Vipp1_C16-C17_ rings contains seven (Fig. 2B).

Within each rung, Vipp1 subunits form a staggered polymer with subunit *j* contacting neighbouring subunits *j*+1 and *j*+3 (Fig. 3B). Hairpin motifs pack side by side so that helices *α*1, *α*2 and *α*3 of subunit *j* form an extended interface underneath the hairpin of neighbouring subunit *j*+1. Concurrently, the helix *α*5 C-terminus (*α*5_C_) of subunit *j* binds across the hairpin tip of subunit *j*+3 forming a contact termed here Interface 1. Importantly, similar hairpin stacking and the equivalent hairpin/helix *α*5 interface is observed within the CHMP1B filament between subunit *j* and subunit *j*+4 (Fig. 3B) (*17, 23*). The hairpin-helix *α*5 contact constitutes a domain swap in CHMP1B when polymerised, and is analogous to the hairpin/helix *α*5 contact observed in other closed ESCRT-III conformations (*16, 31*). Interface 1 is conserved and was predicted for Vipp1 by our co-evolutionary analysis (Fig. 1B and fig. S5). To probe the contribution of Interface 1 in polymer assembly, Vipp1 truncations were generated removing either the helix *α*6 C-terminal extension (Vipp1*Δα*6_1-219_) or helices *α*5 and *α*6 (Vipp1*Δα*5/6_1- 191_). Based on the Vipp1_C11-C17_ models where helix *α*5_C_ is positioned at the tip of the spike (Fig. 2A), *α*6 helices likely coat the ring outside surface and do not contribute directly to polymer formation. Accordingly, both native Vipp1 and Vipp1*Δα*6_1-219_ formed rings and filaments as assayed using gel filtration and negative stain EM. In contrast, Vipp1*Δα*5/6_1-191_ ring and filament formation was abolished (fig. S6B-D). This result showed that helix *α*5 and Interface 1, both of which are conserved in ESCRT-III systems, are essential for Vipp1 filament and ring assembly.

### Conserved hinge regions facilitate domed curvature within rings

The basic building block (or asymmetric unit) of Vipp1_C14_ constitutes a stack of six monomers which when repeated by packing side by side assemble the ring (Fig. 4A). Intriguingly, the six subunits within each asymmetric unit are observed in six distinct conformations. Three hinge regions mediate this conformational versatility (Fig. 4B and movie S1). They are located at the C-terminus of helix *α*2 (Hinge 1 or shoulder), between helix *α*3 and *α*4 (Hinge 2 or elbow), and between helix *α*4 and *α*5 (Hinge 3 or wrist). The tips of the Vipp1 subunit curl progressively upwards if the subunit is positioned in a rung above the ring equator (negative curvature) or downward if positioned below (positive curvature). This curling gives each asymmetric unit a crescent shape, which allows both side by side packing and dome shaped curvature to be achieved. Importantly, the CHMP1B subunit also utilises the three Vipp1 hinges to attain different filament curvatures and to drive helical filament constriction. In particular, Hinge 1 has not previously been reported yet it accounts for a significant degree of the observed flexibility in both Vipp1 and CHMP1B. In both proteins it flexes equivalently (∼6°) albeit along a different axis (Fig. 4B). Overall, the Vipp1_C14_ structure reveals that the underlying principles for how Vipp1 flexes and achieves constriction, albeit with modification, are conserved in CHMP1B and may be usefully applied to better understand other ESCRT-III members as well.

**Fig. 4.**
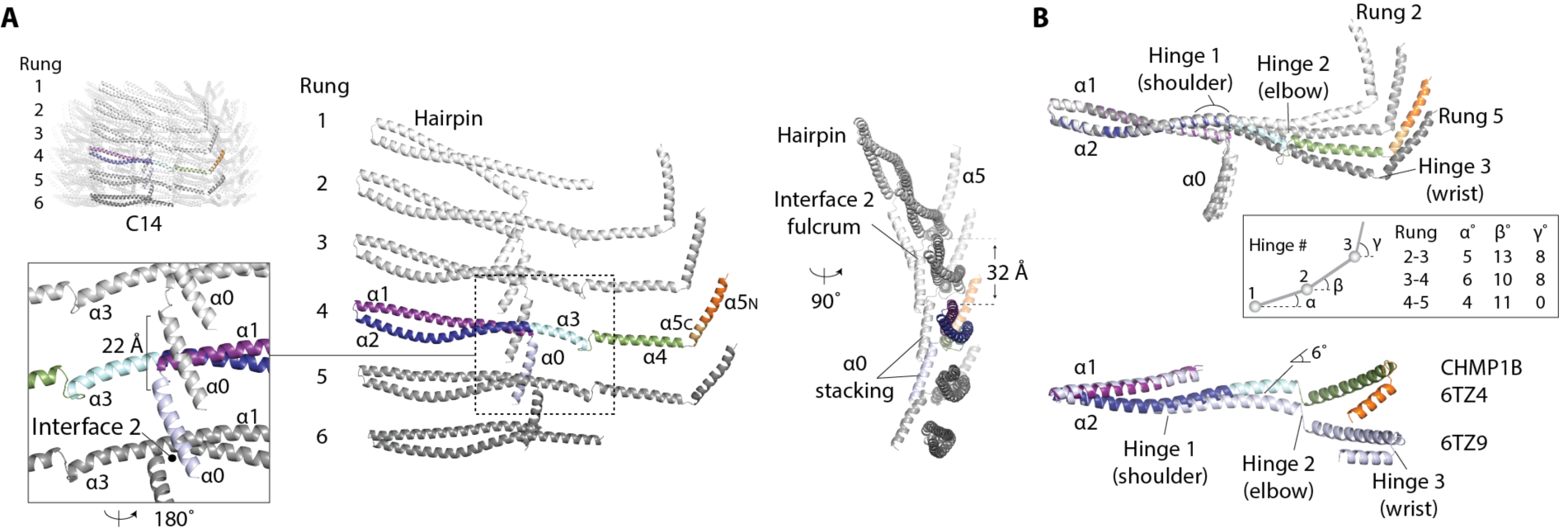
Analysis of the Vipp1_C14_ ring asymmetric unit. (**A**) The ring asymmetric unit comprises an axial stack of six subunits each in its own distinct conformation. Subunits in rung 1 and 6 have disordered helices *α*4-*α*6. Zoom box shows Interface 2 formed from helix *α*0 contacting neighbouring helix *α*0 and helix *α*1 N-terminus. (**B**) Vipp1 and CHMP1B share conserved flexible joints called Hinge 1-3 (shoulder, elbow and wrist). Superposition of Vipp1 asymmetric unit subunits from rungs 2-5 aligned onto hairpin helices *α*1 and *α*2. Subunits transition between negatively and positively curved conformations from the top (rung 2) to the bottom (rung 5).

### How rung stacking induces dome-like curvature

Vipp1 polymers self-organise so that each rung modulates the conformation of all other rungs within the ring stack. Despite the apparent complexity of the Vipp1 ring, each rung associates with its neighbour through just four interfaces. At the Vipp1 N-terminus, helices *α*0 stack axially so that they line the ring lumen (termed Interface 2). In this way, helices *α*0 are well positioned to bind membrane (Fig. 2A, Fig. 4A and fig. S4A). As well as binding rungs together, Interface 2 also serves as a fulcrum by mediating a rotation between each helix *α*0 so that the subunits towards the ring top and base increasingly tilt around the equatorial subunits (Fig. 4A). In this way, the sequential tilting of each helix *α*0 creates the dome shaped curvature of the ring lumen. Interface 3 forms between the N-terminus of helix *α*5 (*α*5_N_) and the hairpin tip from the rung below (Fig. 5A). The amino acids implicated in Interface 3 are highly conserved in both Vipp1 and PspA family members. However, Interface 3 may not form in ESCRT-III as these proteins do not include the helix *α*5 N-terminal extension (Fig. S5). The remaining inter-rung contacts Interface 4 and 5 form smaller packing interfaces between helix *α*4 and helix *α*2, and between Hinge 2 and helix *α*1, respectively (Fig. 5A). Rungs at the bottom of the Vipp1_C14_ ring are tilted relatively flat with helix *α*5 in rung 5 observed 40° from the horizontal (Fig. 5B and fig. S7A). As domed curvature increases towards the top of the ring, helix *α*5 rotates upwards to a maximum of 80° in rung 2. Due to Interfaces 1 and 3, the two hairpins bound to helix *α*5 co-rotate with the effect that they are pulled inward pivoting around Interface 2 (fig. S7B). Thus, we hypothesise that stacking rungs cause lattice tension to build until the capping subunits reach a maximum bending limit. At this point, additional bound subunits cannot flex sufficiently to form Interface 1 or 3, or both, so that partially disordered end rungs form as observed for Vipp1_C14_ rungs 1 and 6.

**Fig. 5.**
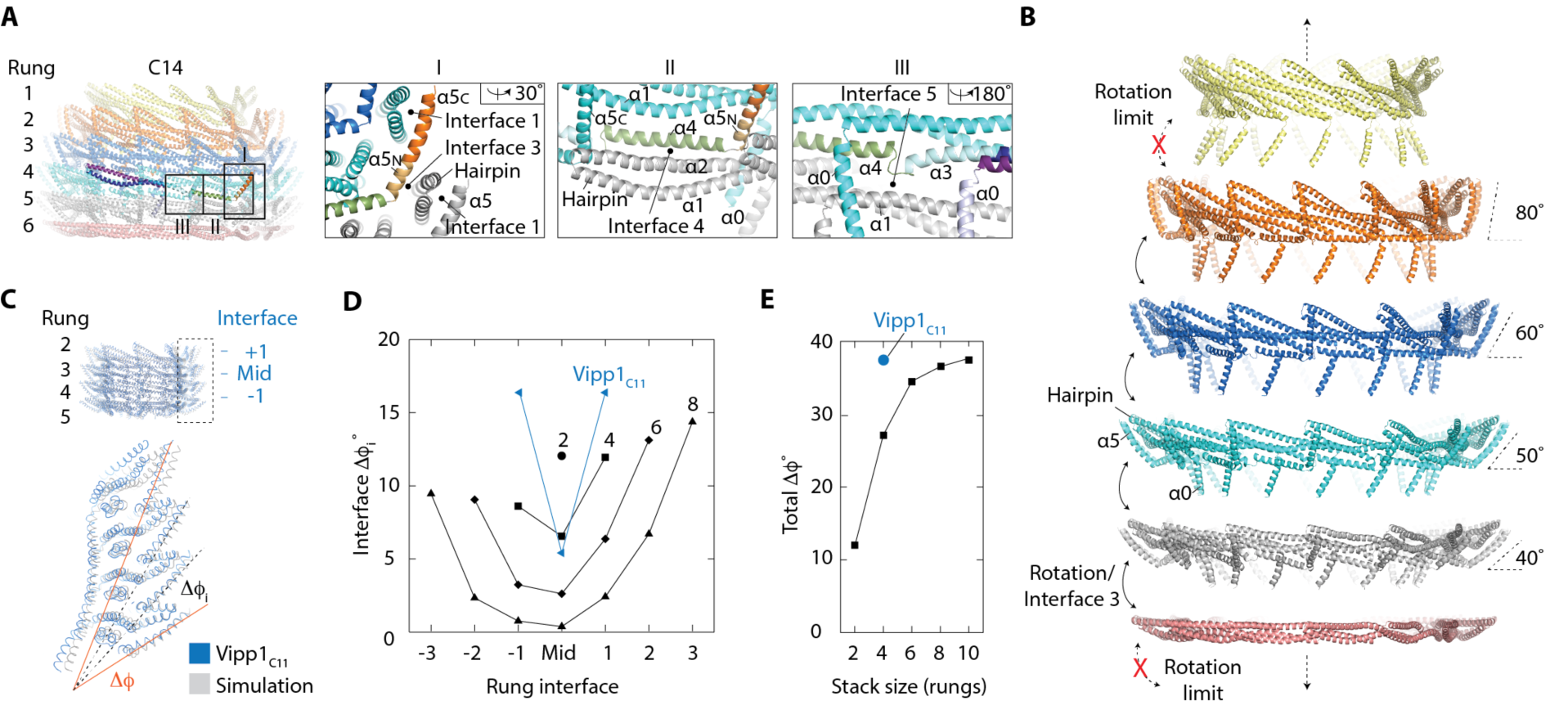
Mechanism for Vipp1 axial or domed curvature. (**A**) Analysis of inter-rung contacts. Zoom boxes show Interfaces 3-5, which combined with Interface 2 (Fig. 4A), define all inter-rung contacts. (**B**) Exploded side view of Vipp1_C14_ shows how each ring comprises a stack of discrete rungs. Each rung constitutes a circular Vipp1 polymer with a distinct conformation. Helix *α*5 is sandwiched between an intra-rung hairpin (Interface 1) and an inter-rung hairpin (Interface 3), which rotate collectively to induce ring constriction and curvature. Geometric constraint ultimately limits constriction. (**C-E**) MD simulations based on Vipp1_C11_ show that inter-rung interfaces define domed curvature. **C** shows overlay of the Vipp1_C11_ structure with a four rung simulation resulting in an equilibrium structure with dome-shaped curvature. *Δ*ϕ_i_ = inter-rung rotation of helix *α*5. *Δ*ϕ= cumulative rotation of helix *α*5 over all rungs. **D** shows *Δ*ϕ_i_ plots from simulations with 2, 4, 6 and 8 rung stacks with C11 symmetry. *Δ*ϕ_i_ for Vipp1_C11_ rungs 2-5 (blue). The largest rotations are observed at the ring top and bottom. **E** shows *Δ*ϕ as a function of rung stack size. Geometric constraints limit the total *Δ*ϕ to near ∼40° regardless of stack size. The limit in *Δ*ϕ from simulated stacks matches the *Δ*ϕ from Vipp1_C11_.

### Modelling dome formation

To test our hypothesis that domed curvature is controlled by a combination of inter-rung contacts and geometric constraints, an idealized elastic-network model of the Vipp1_C11_ ring was constructed. First, an ‘average rung’ was defined that represented the average shape of a single isolated Vipp1_C11_ rung. Then, the way each rung interacts with its neighbouring rungs was defined based on the contacts between rungs 3 and 4 of the Vipp1_C11_ structure (Fig. 2B and Fig. 5C). Finally, Vipp1 rings were initialized as cylindrical stacks of between 2 and 10 average rungs, which were then minimised subject to the intra-rung and inter-rung elastic network (movie S2). The model rings formed Vipp1-like domes with helix *α*5 rotation (*Δ*φ_i_) increasing with rung distance from the ring equator (Fig. 5D). Since, by construction, the model rings had to maintain all their intra- and inter-rung contacts, the strain increased with each additional rung interface. Despite this, the cumulative helix *α*5 rotation (*Δ*φ) between all rung interfaces plateaued near 40°, indicating a strict geometrical bending limit. Remarkably, this maximal rotation within the simulated stacks closely matched the measured experimental *Δ*φ in Vipp1_C11_ (Fig. 5E). This maximal rotation also matched the cumulative ∼40° helix *α*5 rotations observed for Vipp1_C14_ (Fig. 5B). Thus, the simulations suggest a simple self-regulatory mechanism for limiting ring stack size where strain from inter-rung rotation builds up until the formation of Interfaces 1 and 3 is no longer favourable beyond a geometrical limit of ∼40°, after which further stacking is inhibited. Consistent with this, in Vipp1_C11-C17_ rings, Interface 3 was never observed between the top two rungs because the limiting geometry was already reached between the rungs immediately below. To test the importance of Interface 3 for domed curvature formation, we mutated two conserved residues F197K and L200K (Vipp1_F197K/L200K_) within this interface. For the majority of the sample, self-assembly was abolished indicating that Interface 3 was important for both rung formation and ring stability. However, a minor fraction still formed rings and filaments. The filaments were particularly striking as they were unusually long and had a broadly uniform diameter consistent with inter-rung tilt being impeded (fig. S8A). These data suggest that Interface 3 forms a tensile connection between rungs that facilitates the formation of domes. One way to shift ring stack size is to tune geometric stress within the asymmetric unit by changing ring symmetry. Superposition of Vipp1_C11-C17_ asymmetric units showed how subunit bending (movies S3 and S4) varies between Vipp1_C11-C17_ rings. Increasing ring symmetry and consequently ring diameter correlates with geometric stress release within the asymmetric unit. Tipping points occur between Vipp1_C11-C12_ and Vipp1_C15-C16_, which allows rings to increase rung number from five to seven.

### Vipp1 polymers and membrane remodelling

While the structural analysis above focused on domed rings, we also observed native Vipp1 and Vipp1*Δα*6_1-219_ forming multiple types of polymeric assemblies including stacks of rings, ribbons and filaments (fig. S2B, S2D and fig. S8A). Fourier Bessel analysis suggest Vipp1 filaments to be helical with variable diameters and lattices. For the 14 nm diameter filament the dominant layer line corresponds to a 32 Å repeat, which matches the average axial rise between hairpins within neighbouring rungs (fig. S2D and Fig. 4). Given the Vipp1 subunit is > 16 nm in length when maximally curved, significant conformational change or lattice rearrangements may be required to build the 14 nm filament. Stack arrays were observed morphing from discrete rings into ribbons then filaments, particularly with Vipp1*Δα*6_1-219_ (*21*) (fig. S8B). Overall, these micrographs indicate that Vipp1 and Vipp1*Δα*6_1-219_ can transition between lattices in an extraordinary display of assembly versatility reminiscent of ESCRT-III filaments (*4, 17, 32*-*34*).

To determine how Vipp1 might bind and remodel lipids, we first used a spin pelleting assay to show that *N. punctiforme* Vipp1 binds *E. coli* liposomes (fig. S8C). We then directly observed the interplay between Vipp1 and ∼1 µm diameter liposomes by negative stain EM. Vipp1 rings decorated the liposome surface (Fig. 6A and fig. S8D). Side views of Vipp1 bound to liposome edges clearly showed the rings attached to the membrane surface. Tilt and side views also revealed how the ring preferentially attached to the membrane via its bottom rung so that the dome-shaped curvature projected away from the lipid surface. Side views were not observed unless associated with a liposome edge. Stacks of up to six rings were often observed protruding from liposome edges (fig. S8D). Stacks sometimes comprised rings of decreasing diameters to form cones. *N. punctiforme* Vipp1 was also observed tethering liposomes through both single rings and through bridges of stacked rings or cones (Fig. 6A and fig. S8D). Tethered liposomes were consistently attached at the top and bottom of the Vipp1 ring. The observation that liposomes tended not to attach to the side of the ring suggested that helix *α*6, which coats the ring surface and may bind membrane (*21*), was not a key driver of liposome tethering. To further understand how Vipp1 interacts with membrane, we exposed Vipp1 to *E. coli* lipid monolayers (Fig. 6B, 6C and fig. S8E-G). A mosaic of rings was observed bound to the monolayer with end on views only present. Intriguingly, rings bound to lipid monolayers often had occluded lumens demarcated by white centres (Fig. 6C). These occluded rings were consistent with a model where the monolayer was adsorbed onto the inner ring wall forming an encapsulated vesicle-like bud that excludes the negative stain.

**Fig. 6.**
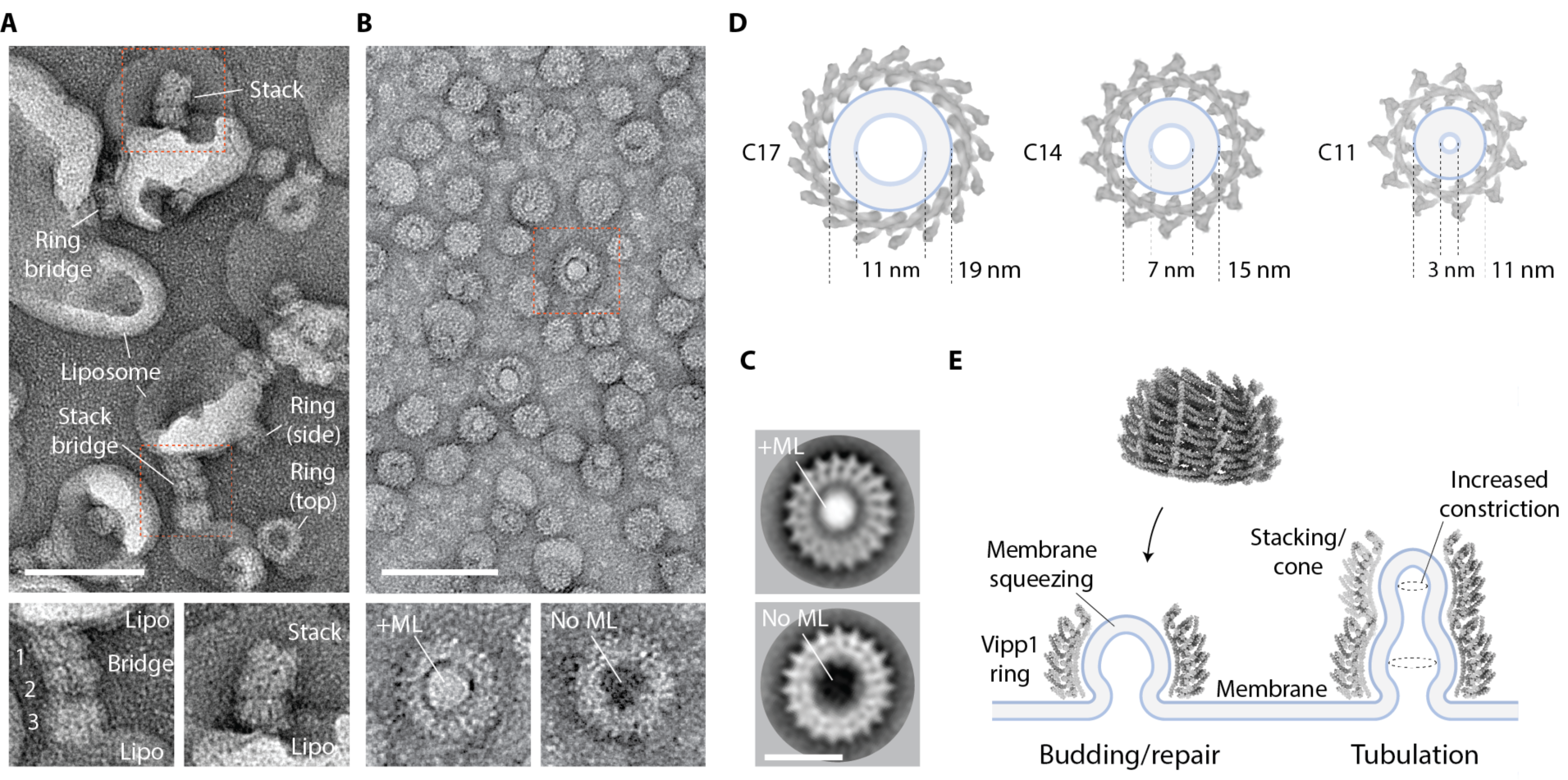
Mechanism of Vipp1 membrane repair and fusion. (**A**) Vipp1 rings decorate and tether liposomes together. Individual rings or ring stacks form bridges between liposomes. Scale bar = 100 nm. (**B**) Vipp1 rings decorate a lipid monolayer (ML). Zoom panels compare rings in the presence or absence of lipid monolayer. Lipid monolayer is drawn into the ring and occludes the lumen (white central density). In the absence of lipid monolayer, rings have an empty lumen (black central density). (**C**) Class averages of Vipp1 rings in the presence (top) and absence (bottom) of lipid monolayer. (**D**) A membrane tube comprising 4 nm lipid bilayer is modelled into Vipp1_C17/C14/C11_ rings to show constriction progression. Membrane hemifusion is expected to be achieved within Vipp1_C11_. (**E**) Schematic showing how Vipp1 may repair damaged or perturbed membrane on a single lipid bilayer. Ring stacks and cones facilitate membrane tubule formation and increasing constriction towards the cone apex.

## Discussion

Here, we show that Vipp1, PspA and ESCRT-III constitute an ancient superfamily of related membrane remodelling proteins conserved across archaea, bacteria and eukaryotes. In addition to homology at the primary amino acid sequence level (fig. S1A), the Vipp1 subunit shares similar secondary structure and overall fold with ESCRT-III proteins (Fig. 3A). When polymerised, Vipp1 shares core assembly features with CHMP1B (*17, 23*). This includes regions of flex that allow both polymers to assume different forms to undertake their function (Fig. 4). Vipp1 contains a similar hairpin motif, elbow, and wrist joint (Hinges 2 and 3) as CHMP1B (*17, 23*). As we show here, both Vipp1 and CHMP1B also have a shoulder joint located at the C-terminus of helix *α*2 (Hinge1). Collectively, these conserved hinges enable the polymers to assume forms that differ widely in curvature and tilt, including a broad variety of complex 3D structures like domes (*35*). In addition, both Vipp1 and CHMP1B form polymers through side-by-side packing of the hairpin motif and through helix *α*5 contacting the hairpin of neighbouring subunits *j*+3 or *j*+4, respectively. This helix *α*5 contact, which forms Interface 1, is conserved and represents a defining feature that explains how this superfamily of proteins generate polymers.

Whilst both Vipp1 and CHMP1B form helical filaments, here we describe the architecture of seven 1.6-3.4 MDa Vipp1 rings from the cyanobacteria *N. punctiforme*. The sequence of ring symmetries presented ranging from C11-C17 allows an unprecedented snapshot into the structural flexibility that enables Vipp1, PspA and ESCRT-III family members to perform their functions. Furthermore, as each ring comprises between 5-7 rungs stacked together to build a unique dome-shaped architecture, our models show the first glimpse into the structural features that enable polymer or filament tilt in an ESCRT-III-like system. This is important as simulations have shown that filament tilt may facilitate ESCRT-III transition from a planar spiral to a three-dimensional cone so as to generate force and drive membrane deformation (*35*).

The physiological role of Vipp1 within the cell remains enigmatic with possible functions in mitigating membrane stress in photosynthetic membranes as well as in thylakoid membrane biogenesis and repair. The closely related bacterial PspA is known to mediate inner membrane repair in response to membrane stress. With only relatively low resolution structural data available for both the Vipp1 and PspA rings (*13, 36*), it is not well understood how Vipp1 and PspA mitigate membrane stress. Here, our Vipp1_C11-C17_ structures suggest a mechanism for stabilising or repairing localised sections of breached or perturbed membrane. They also provide a mechanism for Vipp1 and PspA mediated membrane curving and tubulation. When membrane was modelled into the Vipp1_C11-C17_ structures as a 4 nm bilayer, sequential constriction was observed with the membrane lumen reaching ∼ 3 nm within Vipp1_C11_ (Fig. 6D). This is important as membrane hemifusion or hemifission is achieved when the membrane lumen reaches the approximate thickness of the bilayer (*37, 38*). As we show using membrane binding assays (Fig. 6B, 6C and fig. S8E), in the simplest instance Vipp1 rings will bind to membrane breaches or local bilayer perturbations through exposed *α*0 helices at the ring base. *α*0 helices pack as sub-filaments that line the inner lumen of the ring and form a continuous axial membrane binding interface. This arrangement suggests that a capillary action mechanism draws the membrane into the ring lumen to form a highly curved tubule. It also explains why liposomes are pulled into the central lumen of preformed *Chlamydomonas* Vipp1 helical filaments (*20*). As Vipp1 rings are most constricted at their top, membrane becomes increasingly squeezed as it ascends the inner lumen. The Vipp1_C11_ inner lumen diameter is sufficiently constricted to achieve membrane hemifusion, meaning breached or perturbed membrane leaflets converge to a point of stabilisation or fusion at the dome apex (Fig. 6E). This is consistent with the observation that *Synechocystis* Vipp1 bound to liposomes bends the membrane in regions proximal to the ring (*36*). Additionally, ring stacking and cone formation provide a mechanism to sequentially suck and constrict membrane into tubules (Fig. 6E). Overall, this model is similar to the previously proposed mechanism of ESCRT-III mediated membrane fusion to repair damaged endolysosomes - albeit that in this case membrane remodelling was proposed to occur via the constriction of spiral filaments rather than rings (*39*). Given Vipp1 mediated membrane fusion may occur *in vitro* (*21, 30, 36*), it is possible that rings and ring stacks tether and bridge opposed membranes (Fig. 6A and fig. S8D) as a precursor to membrane fusion. In any bridging event, it may be possible that Vipp1 rings, stacks or filaments extend from both opposing membranes so that a seam of opposing hand is created where the polymers merge, as observed in *Chlamydomonas* Vipp1 protein-lipid tubes (*20*).

In summary, our results unite bacterial, plastidial and archaeal PspA and Vipp1 into a family of bacterial ESCRT-III proteins (bESCRT-III). They show that eukaryotic ESCRT-III (eESCRT-III) and bESCRT-III are ancient membrane remodelling machines that were present in some of the earliest of cells and possibly even our last universal common ancestor (LUCA).

## Acknowledgements

We thank eBIC for cryo-EM data collection support. Paul Simpson for in-house EM support.

## Author contributions

D.S. and B.B. identified the homology between Vipp1/IM30/PspA as ESCRT-III proteins and undertook evolutionary analyses with help from T.W. O.B., J.L. and M.T. generated clones. J.L. processed cryo-EM data and generated reconstructions. J.L. built models with H.L. J.L. purified dynamin. M.T. purified proteins and undertook structure cross-linking validation, helix *α*5 and *α*6 truncation study, Interface 3 mutagenesis all with associated EM studies. S.N. undertook Vipp1 liposome and lipid monolayer EM studies. O.B. purified proteins and undertook cryo-EM studies. J.N. undertook molecular dynamic simulations. H.L. and M.B. supervised O.B. H.L. wrote paper apart from introduction and evolutionary analysis sections by D.S. and B.B. All authors provided manuscript feedback.

## Data and materials availability

3D cryo-EM density maps produced in this study have been deposited in the Electron Microscopy Data Bank with accession code EMD-11468, EMD-11469, EMD-11470, EMD-11478, EMD-11481, EMD-11482 and EMD-11483 for Vipp1_C11-C17_, respectively. Atomic coordinates have been deposited in the Protein Data Bank (PDB) under accession code 6ZVR, 6ZVS, 6ZVT, 6ZW4, 6ZW5, 6ZW6 and 6ZW7 for Vipp1_C11-C17_, respectively.

## Supplementary Figures

**Fig. S1.**
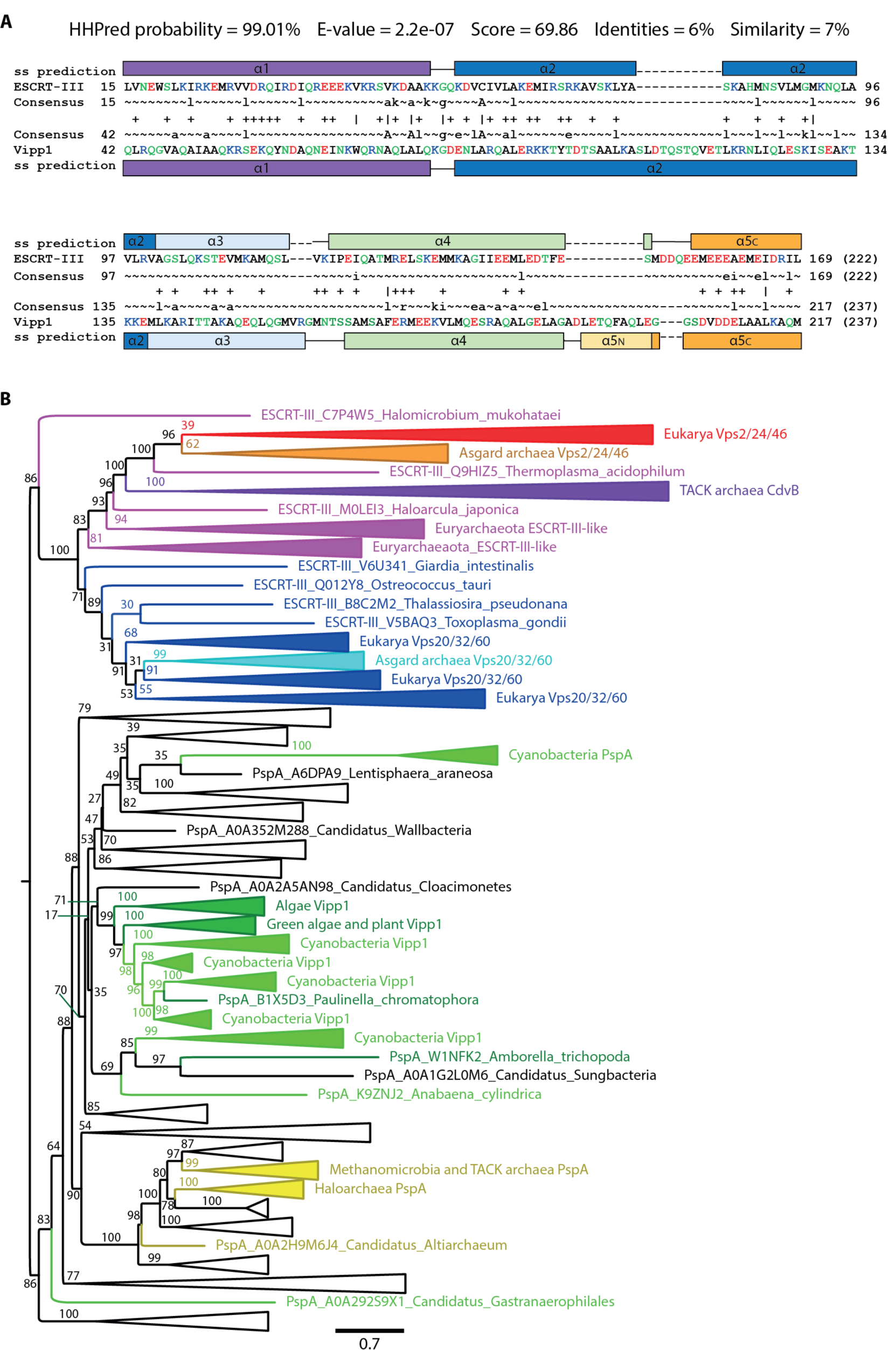
Evolutionary relationship of ESCRT-III/Snf7/CdvB and PspA/Vipp1 proteins. (**A**) ESCRT-III/Snf7/CdvB and PspA/Vipp1 protein families are homologues based on an analysis performed using HHPred (*8*). The alignment of human CHMP3 (ESCRT-III) and *N. punctiforme* Vipp1 is based on their HMM profiles. Only highly conserved (uppercase) and moderately conserved (lowercase) HMM consensus positions are displayed. Vertical lines and plus signs indicate identical and similar HMM positions, respectively. CHMP3 and Vipp1 residues are colour-coded according to their chemical properties: polar – green; positively charged – blue; negatively charged – red; and hydrophobic - black. Secondary structure (ss) predictions are shown with helices numbered based on both CHMP3 (PDB code 3FRT) and CHMP1B (PDB code 3JC1) structures. (**B**) A phylogenetic tree presenting the relationship of the PspA and ESCRT-III families. Unlabelled white collapsed branches contain bacterial PspA homologues. Numbers indicate the relative support from 100 bootstrap replicates. Scale bar represents expected substitutions per site.

**Fig. S2.**
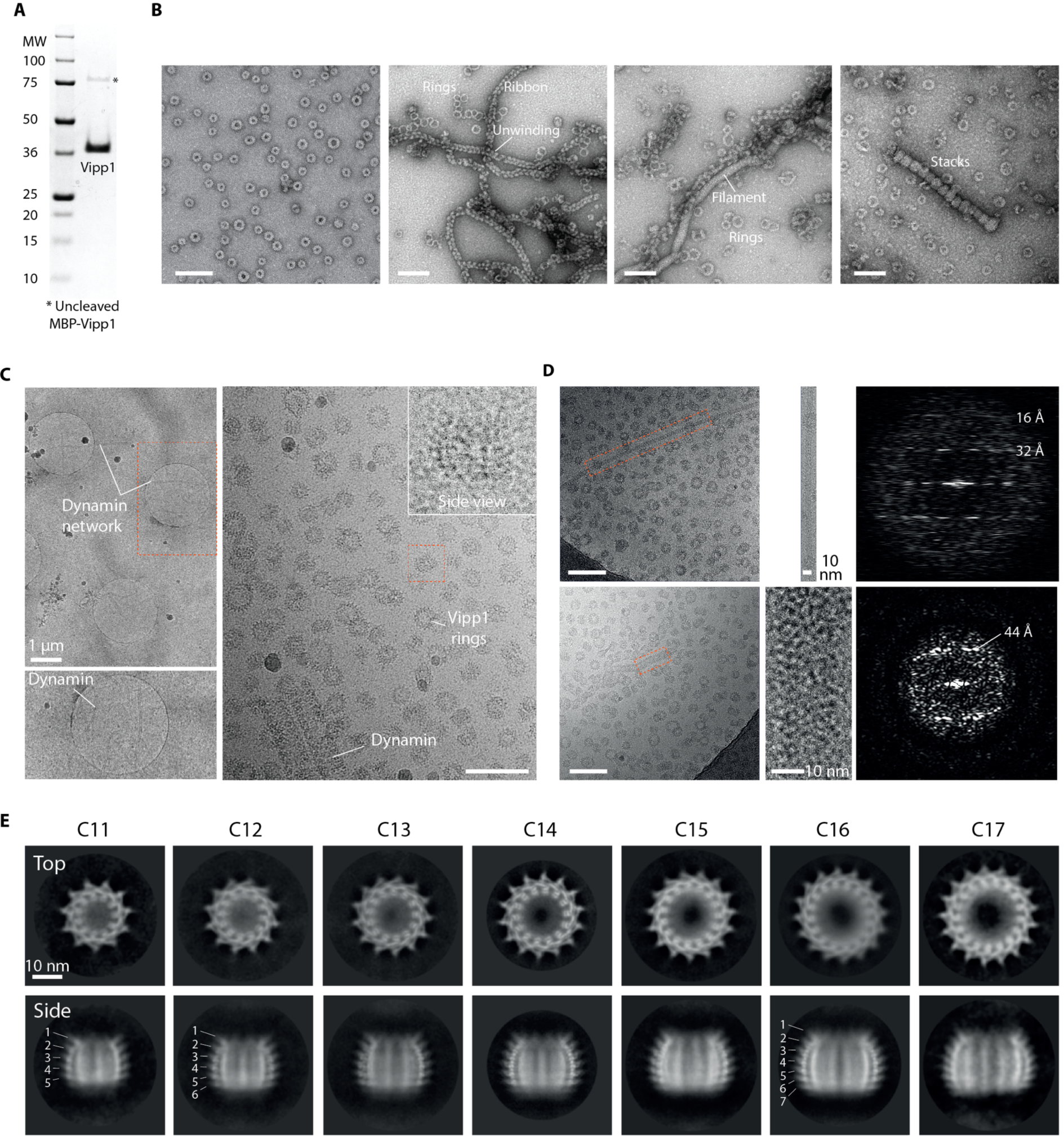
Purification and electron microscopy of *N. punctiforme* Vipp1. (**A**) SDS-PAGE showing purified Vipp1. Note that Vipp1 migrates at ∼38 kDa whereas its true mass is 28.7 kDa. (**B**) Gallery of negative stain EM micrographs showing Vipp1 forming rings, helical ribbons, filaments and stacks. Scale bar = 100 nm. (**C**) Cryo-EM micrographs showing Vipp1 rings mixed with human Dynamin 1 filaments. Left panels-low magnification overview of the holey grid showing the Dynamin 1 filament network used to maintain ice thickness and promote Vipp1 side views. Red rectangle outlines zoom panel below. Right panel-example micrograph showing Vipp1 rings including side views together with the Dynamin 1 filaments. Scale bar = 100 nm. (**D**) Cryo-EM micrographs of selected Vipp1 filaments with associated Fourier Bessel analysis. Vipp1 filaments were sometimes observed amongst Vipp1 rings. A 14 nm filament (top) shows a helical repeat at 32 Å and 16 Å. 32 Å matches with the axial rise between the hairpin motif of neighbouring rungs. A 22 nm diameter filament (bottom) shows an axial repeat at 44 Å. Scale bar = 100 nm. (**E**) Gallery of Vipp1_C11-C17_ 2D class averages from the cryo-EM dataset showing end and side views.

**Fig. S3.**
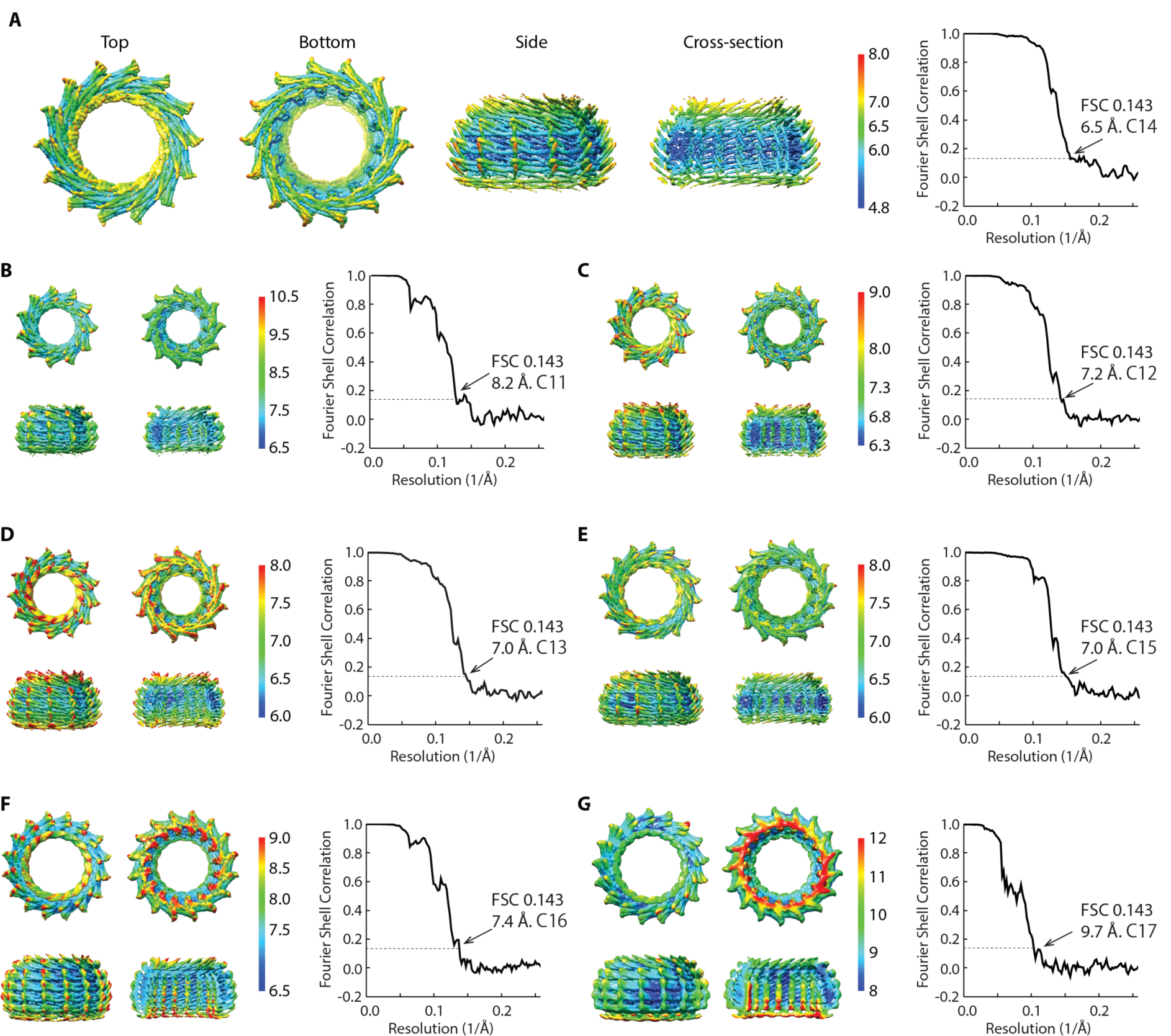
Vipp1_C11-C17_ local resolution map and FSC curves. (**A-G**) Gallery of sharpened maps contoured between 4-6*σ* showing local resolution estimates and associated gold standard FSC curves.

**Fig. S4.**
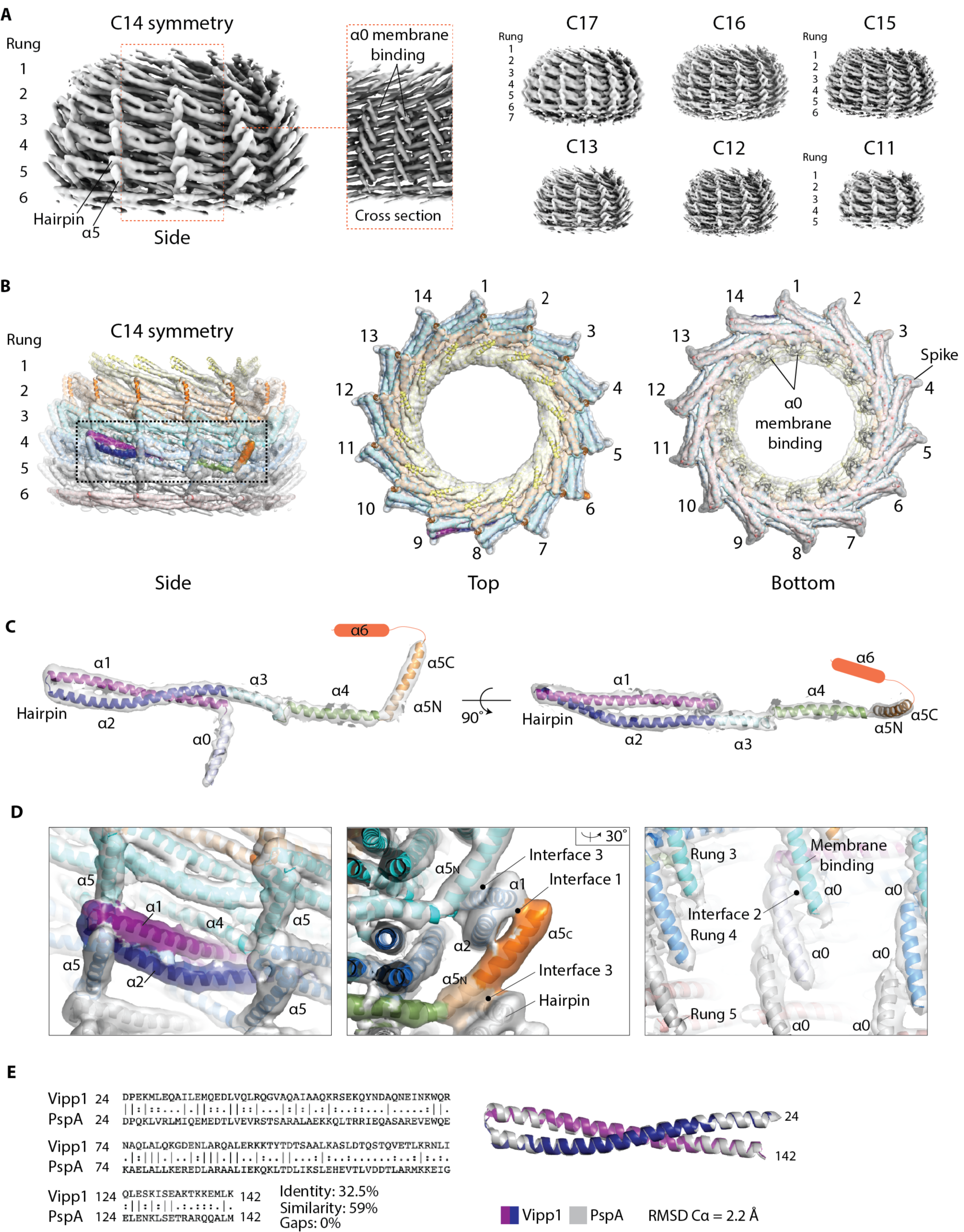
Vipp1_C11-C17_ map quality and model building. (**A**) Gallery of Vipp1_C11-C17_ EM density maps contoured between 4-6*σ*. (**B**) Vipp1_C14_ map fitted with 84 Vipp1 subunits. Subunits in rungs 1 and 6 were partially built with helices *α*4, *α*5 and *α*6 omitted due to disordered or absent density. Density for helix *α*6 was never observed. Map contoured at 4*σ*. (**C**) Isolated Vipp1 monomer showing quality of map, build and fit. The subunit extracted is indicated in the rectangular box in **B**. (**D**) Select regions of Vipp1_C14_ showing quality of map, build and fit. Left panel-hairpin motif. Middle panel-end view of a hairpin motif forming both intra-rung Interface 1 and inter-rung Interface 3 with helix *α*5. Right panel-helix *α*0 stacking forms Interface 2. (**E**) Left panel-pairwise alignment of *Nostoc punctiforme* Vipp1 (Uniprot code B2J6D9) with *Escherichia coli* PspA (Uniprot code P0AFM6) amino acids 24-142 (hairpin motif). Right panel-superposition of Vipp1_C14_ amino acids 24-142 (rung number 4) with PspA partial structure (PDB 4WHE). RMSD = relative mean square deviation.

**Fig. S5.**
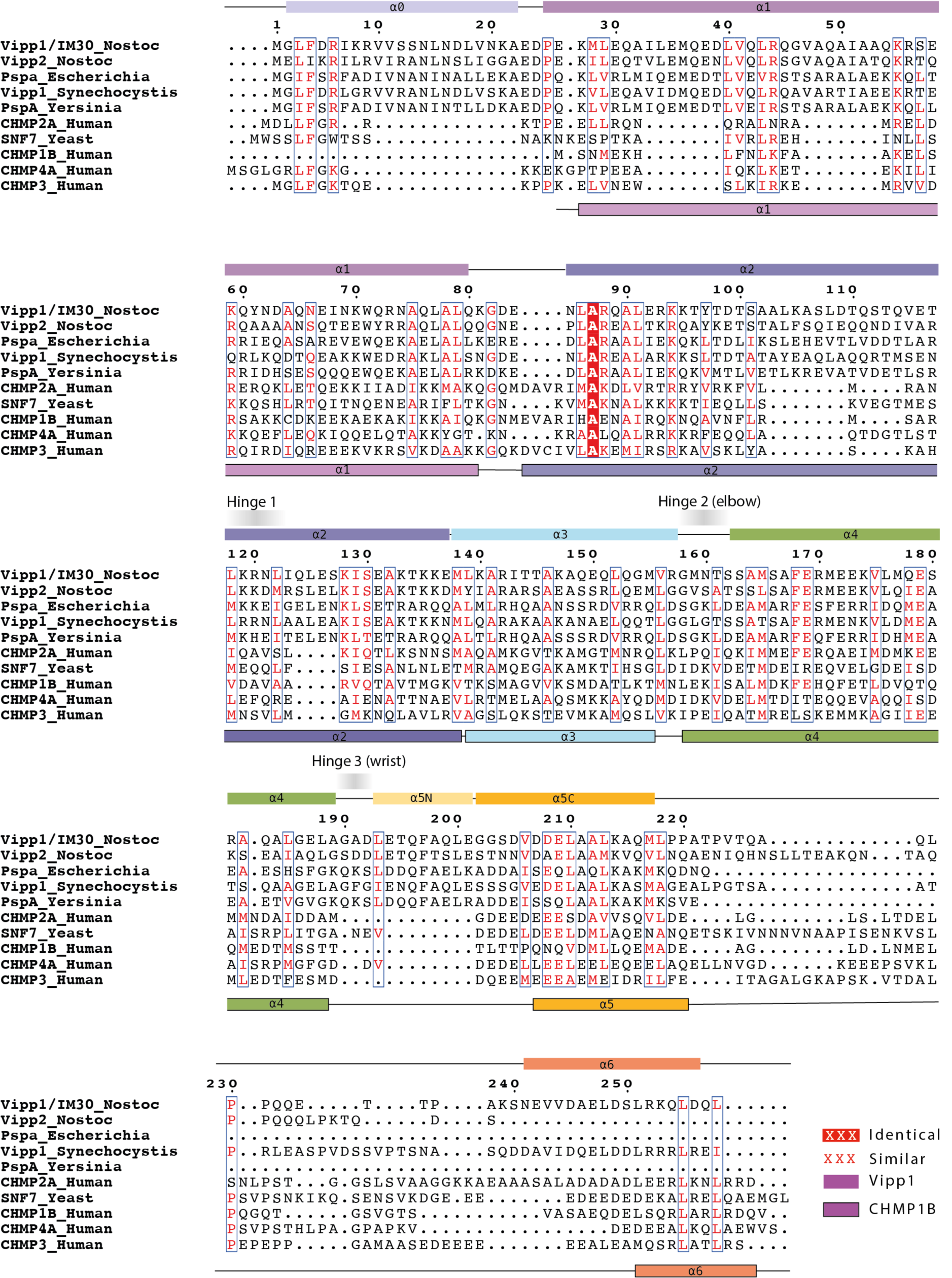
Vipp1 secondary structure assignment and sequence alignment with PspA and ESCRT-III. Sequences were aligned using Clustal Omega and include Vipp1/IM30 *Nostoc punctiforme* (Uniprot code B2J6D9), Vipp2 *Nostoc punctiforme* (Uniprot code B2J6E0), *Escherichia coli* PspA (Uniprot code P0AFM6), *Synechocystis sp*. Vipp1 (Uniprot code A0A068MW27), *Yersinia pestis* PspA (Uniprot code Q0WEH0), human CHMP2A (Uniprot code O43633), yeast SNF7 (Uniprot code P39929), human CHMP1B (Uniprot code Q7LBR1), human CHMP4A (Uniprot code Q9BY43) and human CHMP3 (Uniprot code Q0Y3E7). Cartoon shows Vipp1 (top) and CHMP1B (bottom, PDB code 6TZ4) secondary structure.

**Fig. S6.**
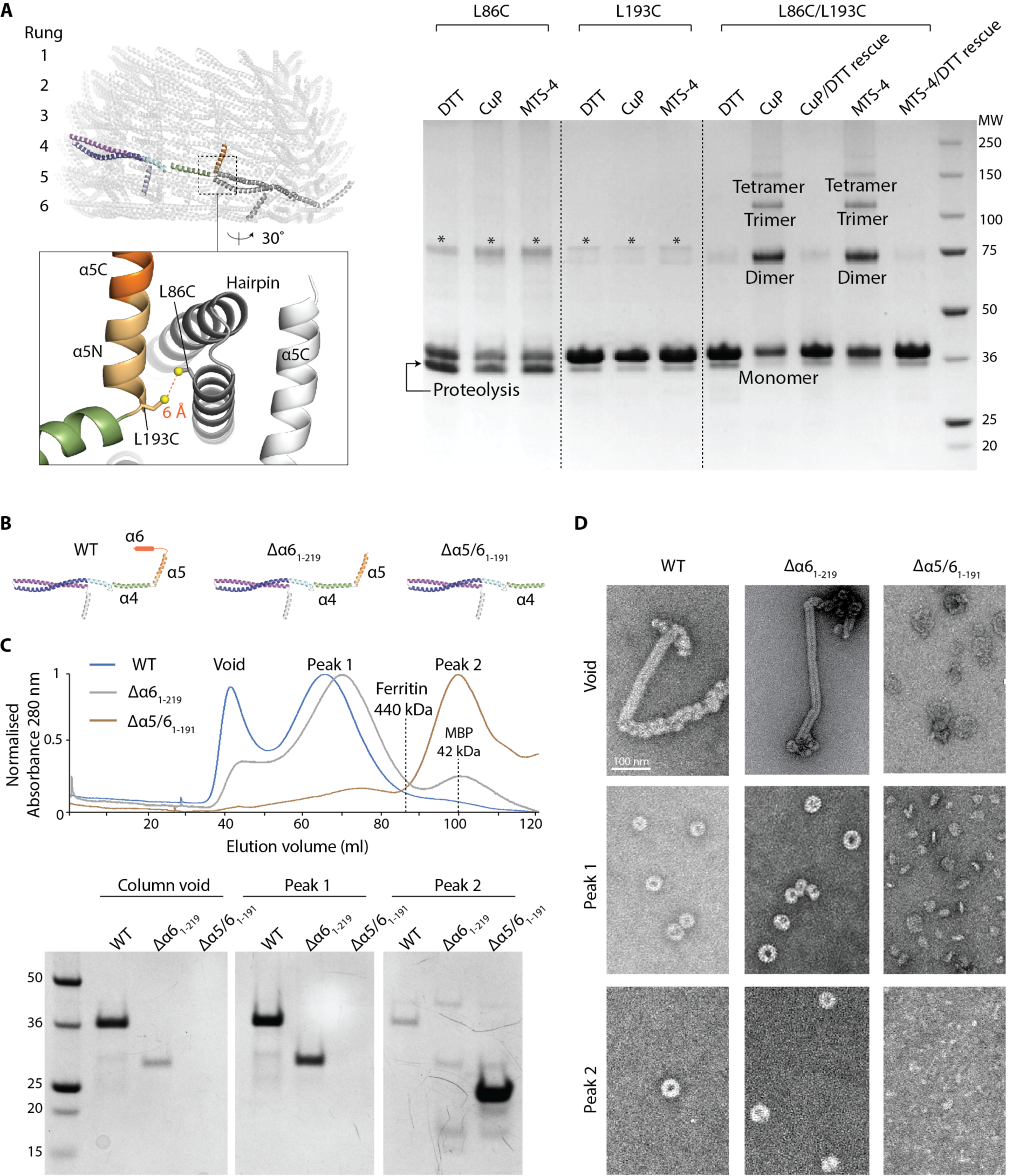
Filament assembly and structure validation. (**A**) Vipp1 monomer structure sequence register validation based on cysteine cross-linking. Zoom box indicates position of L193C and L86C mutants (Vipp1_L86C/L193C_) with the sulphur atoms predicted to be ∼6 Å within the Vipp1_C14_ structure. Vipp1_L86C/L193C_ forms an inter-rung connection between helix *α*5 and the hairpin motif (Interface 3). Disulphide bond formation is observed only in the presence of oxidising agent Cu(II)-phenanthroline (CuP) and MTS4 cross-linker. Disulphide bond formation can be rescued upon subsequent DTT incubation. * indicates uncleaved MBP-Vipp1 fusion protein. Vipp1_L86C_ showed higher levels of proteolysis and resistance to full MBP cleavage than normal presumably due to the sensitive position of the mutation. (**B**) Vipp1 subunit cartoon schematics showing wild type (WT), Vipp1*Δα*6_1-219_ and Vipp1*Δα*5/6_1-191_ secondary structure. (**C, D**) Filament assembly assay. In **C**, comparison of WT, Vipp1*Δα*6_1-219_ and Vipp1*Δα*5/6_1-191_ size exclusion profiles using a sephacryl S-500 resin. Associated SDS-PAGE is shown analysing the column exclusion limit (void volume), Peak 1 containing high molecular weight species greater than Ferritin (440 kDa) and usually associated with Vipp1_C11- C17_ rings, and Peak 2 associated with low molecular weight proteins such as maltose-binding protein (42 kDa). Negative stain EM analysis is shown in **D** for void volume, Peak 1 and Peak 2. WT and Vipp1*Δα*6_1-219_ behave similarly with filaments observed in the void volume and Vipp1_C11-C17_ rings observed in Peak 1. In contrast, Vipp1*Δα*5/6_1-191_ does not form filaments or rings and instead runs as a low molecular weight species consistent with monomer or dimer. This experiment shows that helix *α*5 is essential for polymer formation whilst helix *α*6 is not. It also broadly validates the helix *α*5 and *α*6 domain assignment within the Vipp1 structures.

**Fig. S7.**
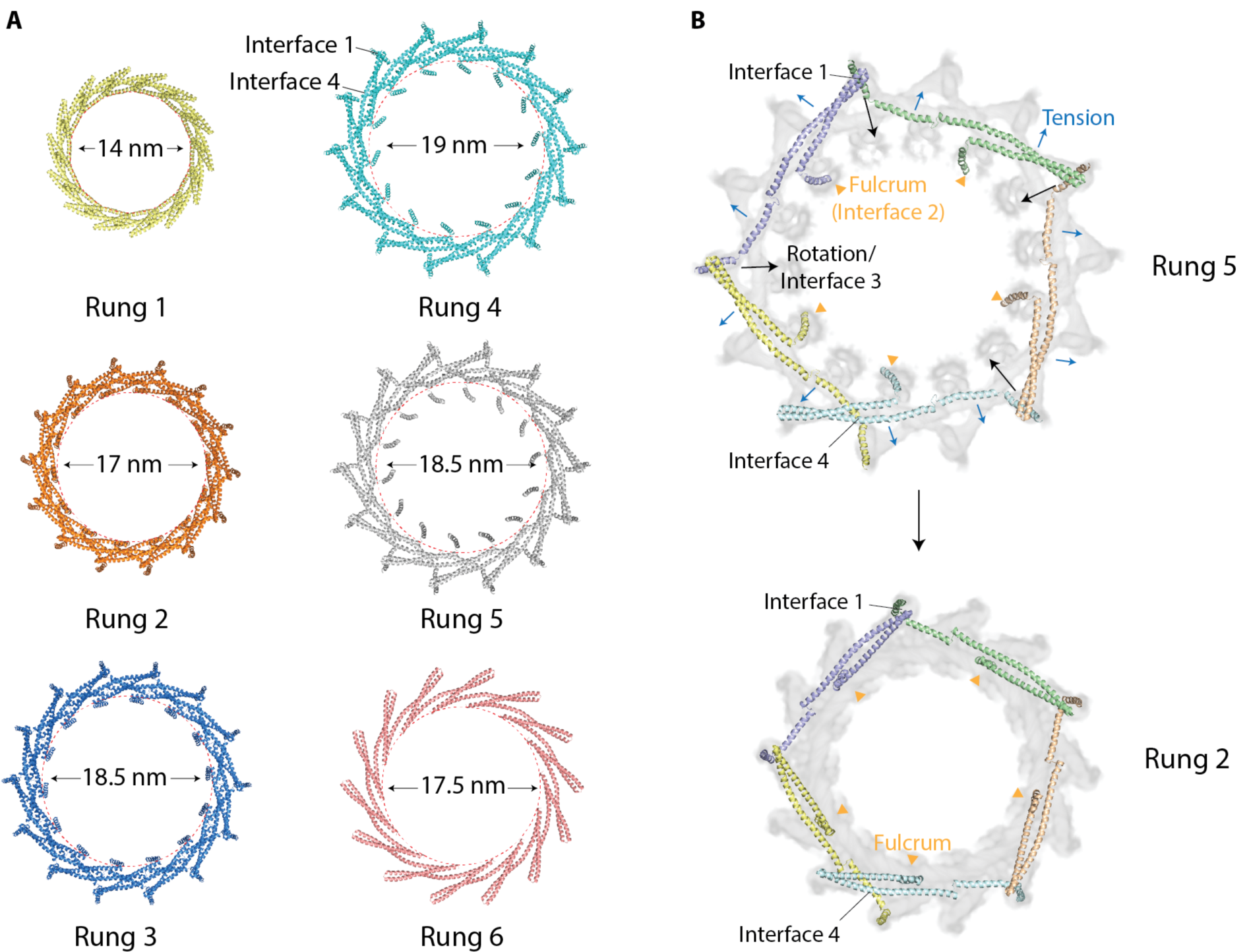
Mechanism for Vipp1 axial or domed curvature. (**A**) Exploded top view of Vipp1_C14_ shows how each ring comprises a stack of discrete rungs. Each rung constitutes a circular Vipp1 polymer with a distinct conformation. Ring diameters were measured at the helix *α*1 N-terminus so as to highlight hairpin constriction between rungs. (**B**) Comparison of rung 5 and rung 2 to highlight the conformational changes required to drive domed curvature. For each rung only those subunits are shown (*j* and *j*+3) that interconnect via Interface 1 to form one turn. Polymer tilt involves hairpin and helix *α*5 rotation (Interface 1 and 3) around Interface 2, which appears to represent a fulcrum within the centre of each subunit. Helix *α*0 within Interface 2 also rotates to create the domed curvature of the inner lumen. Increasing polymer or filament tilt and rotation builds subunit tension until ultimately geometric constraint limits the formation of Interface 3 and/or Interface 1 along with further rung stacking and constriction.

**Fig. S8.**
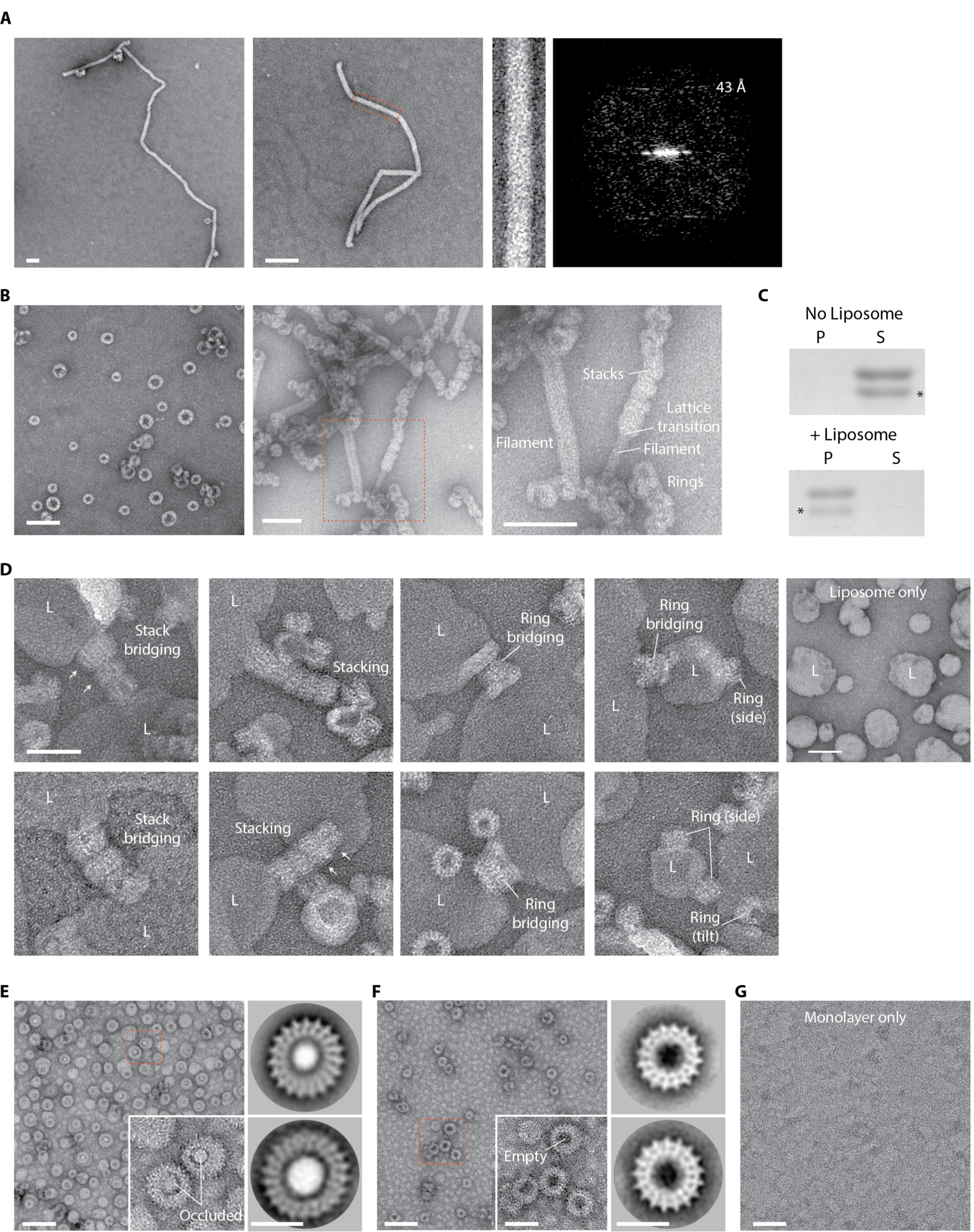
Negative stain EM analyses of Vipp1 and Vipp1 mutants in the presence and absence of lipids. (**A**) Vipp1_F197K/L200K_ forms unusually long filaments with broadly uniform diameter suggesting that loss of Interface 3 impedes filament tilt and axial curving (left two panels). Zoom panel shows close up of the filament outlined by the dotted rectangle. Filament diameter is ∼22 nm. Right panel shows Fourier Bessel analysis of the zoomed filament with a dominant repeat at 43 Å. The filament diameter and likely helical repeat appears consistent with the ∼22 nm Vipp1 wild type filament (fig. S2D). Scale bar = 50 nm. (**B**) Vipp1*Δα*6_1-219_ forms individual rings, ring stacks and filaments. Compared to native Vipp1, Vipp1*Δα*6_1-219_ has a much greater propensity to form ring stacks and filaments. Zoom panel shows region outlined by dotted rectangle where ring stacks can be observed morphing into filaments and acting as nucleation points for filament formation and lattice transitions. Scale bar = 50 nm. (**C**) Native Vipp1 spin assay in the presence and absence of *E. coli* liposomes. Vipp1 is observed in the pellet fraction only in the presence of the liposomes, indicative of lipid binding. * indicates Vipp1 proteolysis. (**D**) Gallery of micrographs showing how Vipp1 rings decorate and tether liposomes (L) together. Individual rings or ring stacks form bridges between liposomes. Vipp1 is rarely observed unattached to a liposome and resting on the carbon surface. Vipp1 stacks may reduce in diameter to form cones (white arrows). Scale bar = 50 nm. (**E**) Left panel - Vipp1 rings decorate a lipid monolayer (ML). Zoom panel shows region outlined by dotted rectangle. Monolayer is drawn into the ring and occludes the lumen (white central density). Right panels-example class averages. (**F)** In the absence of monolayers, Vipp1 rings have empty lumens (black central density). (**G**) Monolayers only. Scale bar = 100 nm.

**Table S1.**
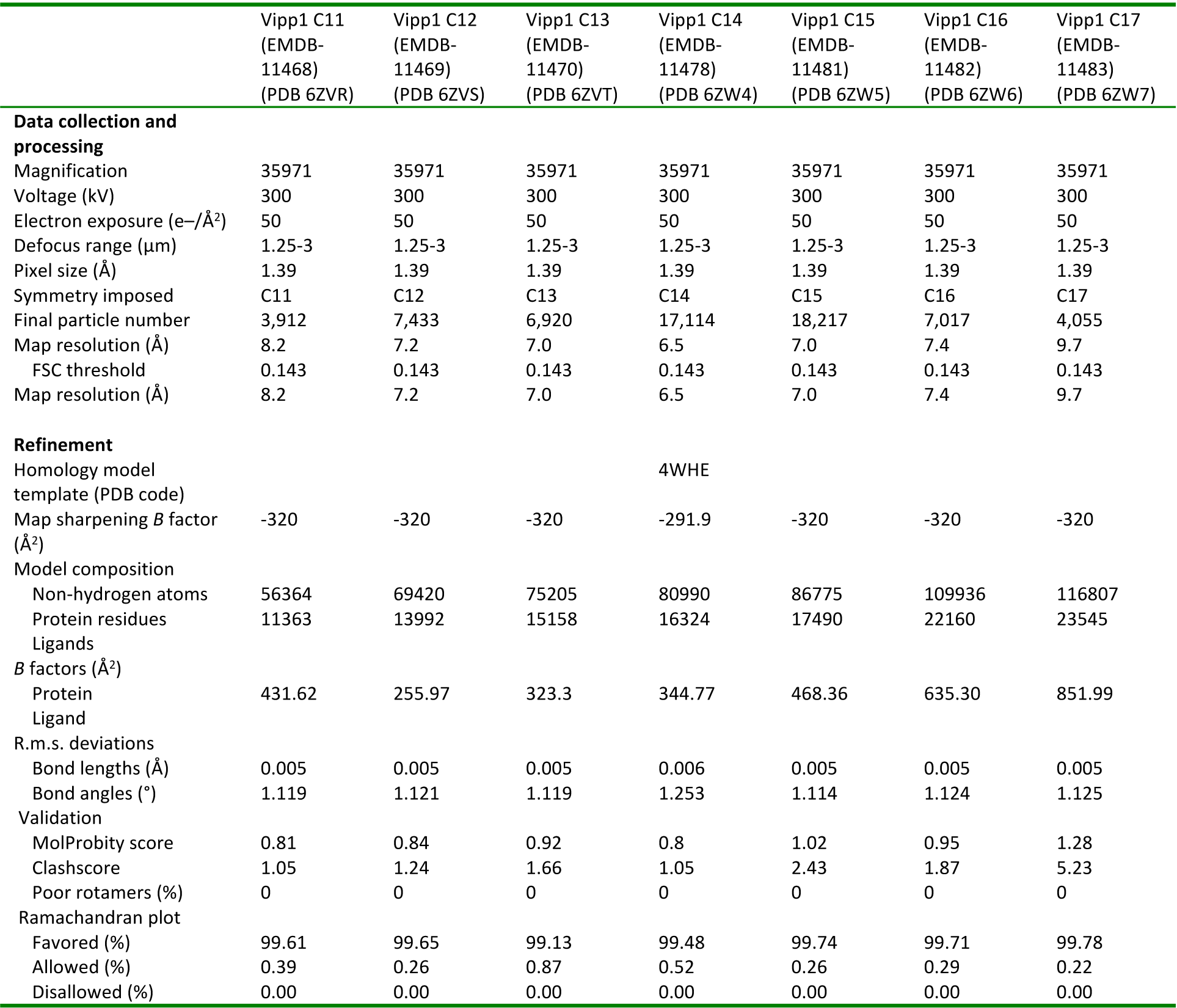
Cryo-EM data collection, refinement and validation statistics.

|  | Vipp1 C11<br>(EMDB-11468)<br>(PDB 6ZVR) | Vipp1 C12<br>(EMDB-11469)<br>(PDB 6ZVS) | Vipp1 C13<br>(EMDB-11470)<br>(PDB 6ZVT) | Vipp1 C14<br>(EMDB-11478)<br>(PDB 6ZW4) | Vipp1 C15<br>(EMDB-11481)<br>(PDB 6ZW5) | Vipp1 C16<br>(EMDB-11482)<br>(PDB 6ZW6) | Vipp1 C17<br>(EMDB-11483)<br>(PDB 6ZW7) |
| --- | --- | --- | --- | --- | --- | --- | --- |
| **Data collection and processing** |  |  |  |  |  |  |  |
| Magnification | 35971 | 35971 | 35971 | 35971 | 35971 | 35971 | 35971 |
| Voltage (kV) | 300 | 300 | 300 | 300 | 300 | 300 | 300 |
| Electron exposure (e<sup>-</sup>/Å<sup>2</sup>) | 50 | 50 | 50 | 50 | 50 | 50 | 50 |
| Defocus range (μm) | 1.25-3 | 1.25-3 | 1.25-3 | 1.25-3 | 1.25-3 | 1.25-3 | 1.25-3 |
| Pixel size (Å) | 1.39 | 1.39 | 1.39 | 1.39 | 1.39 | 1.39 | 1.39 |
| Symmetry imposed | C11 | C12 | C13 | C14 | C15 | C16 | C17 |
| Final particle number | 3,912 | 7,433 | 6,920 | 17,114 | 18,217 | 7,017 | 4,055 |
| Map resolution (Å) | 8.2 | 7.2 | 7.0 | 6.5 | 7.0 | 7.4 | 9.7 |
| FSC threshold | 0.143 | 0.143 | 0.143 | 0.143 | 0.143 | 0.143 | 0.143 |
| Map resolution (Å) | 8.2 | 7.2 | 7.0 | 6.5 | 7.0 | 7.4 | 9.7 |
| **Refinement** |  |  |  |  |  |  |  |
| Homology model template (PDB code) |  |  |  | 4WHE |  |  |  |
| Map sharpening *B* factor (Å<sup>2</sup>) | -320 | -320 | -320 | -291.9 | -320 | -320 | -320 |
| **Model composition** |  |  |  |  |  |  |  |
| Non-hydrogen atoms | 56364 | 69420 | 75205 | 80990 | 86775 | 109936 | 116807 |
| Protein residues | 11363 | 13992 | 15158 | 16324 | 17490 | 22160 | 23545 |
| Ligands |  |  |  |  |  |  |  |
| ***B* factors (Å<sup>2</sup>)** |  |  |  |  |  |  |  |
| Protein | 431.62 | 255.97 | 323.3 | 344.77 | 468.36 | 635.30 | 851.99 |
| Ligand |  |  |  |  |  |  |  |
| **R.m.s. deviations** |  |  |  |  |  |  |  |
| Bond lengths (Å) | 0.005 | 0.005 | 0.005 | 0.006 | 0.005 | 0.005 | 0.005 |
| Bond angles (°) | 1.119 | 1.121 | 1.119 | 1.253 | 1.114 | 1.124 | 1.125 |
| **Validation** |  |  |  |  |  |  |  |
| MolProbity score | 0.81 | 0.84 | 0.92 | 0.8 | 1.02 | 0.95 | 1.28 |
| Clashscore | 1.05 | 1.24 | 1.66 | 1.05 | 2.43 | 1.87 | 5.23 |
| Poor rotamers (%) | 0 | 0 | 0 | 0 | 0 | 0 | 0 |
| **Ramachandran plot** |  |  |  |  |  |  |  |
| Favored (%) | 99.61 | 99.65 | 99.13 | 99.48 | 99.74 | 99.71 | 99.78 |
| Allowed (%) | 0.39 | 0.26 | 0.87 | 0.52 | 0.26 | 0.29 | 0.22 |
| Disallowed (%) | 0.00 | 0.00 | 0.00 | 0.00 | 0.00 | 0.00 | 0.00 |

**Table S2.**
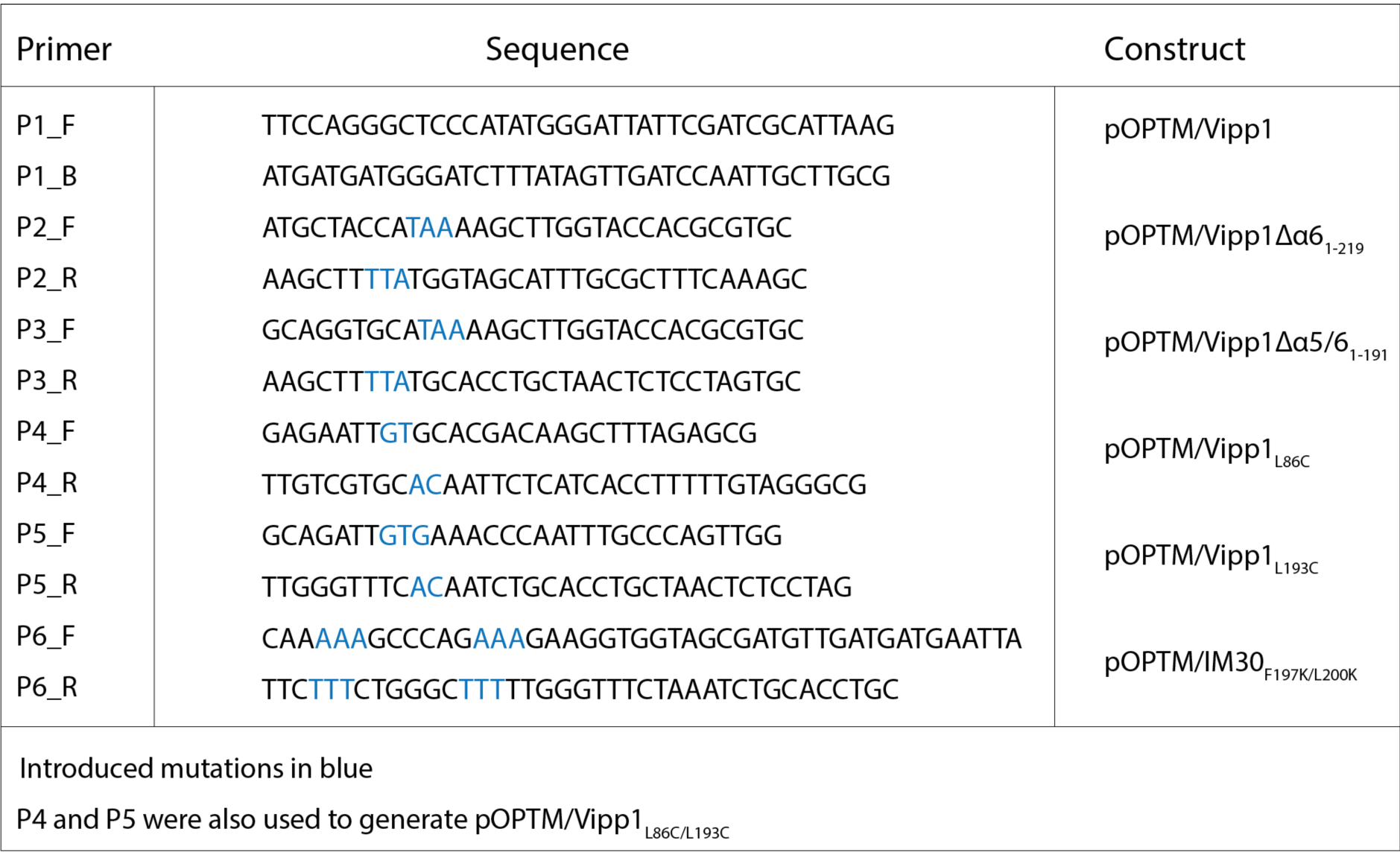
Primers and plasmids.

## Supplementary movies

**Movie S1**. Morph between superposed Vipp1 subunits within the Vipp1_C14_ asymmetric unit. Complete subunits from rungs 2-5 only are shown. Conformational change is mediated by Hinges 1-3 conserved between Vipp1 and ESCRT-III-like proteins.

**Movie S2**. Minimised elastic-network model of Vipp1 forms Vipp1-like domes. The initial structure is a C11 stack composed of eight identical and energy minimized rungs that contains no inter-rung interactions. Inter-rung interactions are added and the movie traces the minimisation of the elastic energy. The equilibrium structure reached is a domed structure that balances the inter-rung rotation favoured by the inter-rung interactions with the overall distortion in each rung.

**Movies S3 and S4**. Front and side view morph between superposed Vipp1_C11-C17_ asymmetric units. Morph shows range of conformational changes between different ring symmetries.

## Materials and Methods

### Methods

#### ESCRT-III homology searches, secondary structure prediction and co-evolutionary analysis

To search for ESCRT-III relatives, we used sensitive protein sequence searches based on Hidden Markov Models (HMMs; HHMER (*40*), HHPred (*8*) and Pfam (*41*)). Several individual eukaryotic ESCRT-III proteins, multiple sequence alignments and HMM profiles were used as queries in these searches. These analyses consistently identified PspA/Vipp1 proteins as the only ESCRT-III homologues in sequence, domain and structural databases (Uniprot (*42*), GenBank (*43*), Pfam (*41*) and PDB (*44*)). This observation was corroborated by the fact that these protein families share a common Pfam clan which only includes PspA and ESCRT-III (Pfam CL0235). Secondary structure predictions were performed using HHPred (*8*), Phyre2 (*45*) and Psipred (*46*) software. To obtain a statistical model of the PspA/Vipp1 family that captures patterns of residue co-evolution, Gremlin (*14, 15*) was used. For this analysis, a total of 2844 homologous proteins were obtained using *N. punctiforme* Vipp1 as a query, four iterations of JackHHMER searches, E-value ≤ 1e-10, using a coverage filter of 50% and gap removal of 75%.

#### Phylogenetic studies

The phylogenetic distribution of the PspA/Vipp1 and ESCRT-III families was generated using AnnoTree (*47*) searches in Pfam (*41*) and an E-value ≤ 1e-05. Homologues of these protein families were retrieved from Uniprot (*42*) by HMMER (*40*) searching and from InterPro (*48*) database, followed by manual inspection. Eukaryotic, archaeal and bacterial PspA and ESCRT-III proteins were selected to achieve a broad distribution of homologues across the tree of life. In total, 264 PspA and 332 ESCRT-III sequences were selected, aligned using the l-INS-I mode in mafft 7.3.1 (*49*), and poorly-aligned regions were identified and removed using the “gappyout” mode in trimAl 1.3 (*50*). The phylogeny was inferred in IQ-Tree 1.6.10 (*51*) under the LG+C30+G+F model, which was the best-fitting model according to the BIC criterion. This model accounts for differences in exchange rates among amino acids (LG), different site compositions (C30+F), and models across-site rate variation using a discretized Gamma distribution with 4 rate categories. Branch supports are ultrafast (UFBoot2 (*52*)) bootstraps.

#### Vipp1 cloning, protein expression and purification

Plasmid mutagenesis and all clones were generated using the Gibson isothermal DNA assembly protocol (*53*). Plasmids and primers used in this study are listed in Table S2. The coding sequence for *Nostoc punctiforme vipp1* (Uniprot code B2J6D9) was cloned into pOPTM (a pET derivative) to yield an N-terminal MBP fusion with a TEV cleavage site in the linker. An N-terminal hexahistidine tag was included on the MBP moiety. For the purification of both native and mutant Vipp1, clones were co-transformed into *E. coli* C43 (DE3) electro-competent cells (Lucigen) modified here to incorporate a *pspA* gene knockout using a Lambda Red recombinase strategy (*54*). Cells were grown on selective LB-agarose plates with ampicillin (100 μg/ml). 2xYT media was inoculated and cells grown at 37 °C until induction at OD600 = 0.8 with 1 mM isopropyl β-D-1-thiogalactopyranoside (IPTG). Cells were grown for ∼15 h at 18 °C and shaken at 220 rpm. All the further steps were carried out at 4 °C unless otherwise specified. Purification of Vipp1_L86C_, Vipp1_L193C_ and Vipp1_L86C/L193C_ were performed in the presence of 2 mM dithiothreitol (DTT). Pellets were re-suspended in ice-cold buffer 50 mM Tris-HCl pH 7.5, 300 mM NaCl, treated with 2 mM MgCl_2_, 0.1 mg/ml DNase I, 0.5 mg/ml lysozyme and sonicated on ice. The lysate was clarified by centrifugation at 16,000 × g for 20 min. The supernatant was incubated with gentle shaking for 1 h with 10 mL of amylose resin (NEB) pre-equilibrated in 50 mM Tris-HCl pH 7.5, 300 mM NaCl (wash buffer). The resin was washed with 100 mL and purified MBP-Vipp1 eluted with wash buffer supplemented with 15 mM maltose. The sample was incubated for 24 h at room temperature with TEV and then dialysed (12-14 kDa MW cut-off) overnight in 25 mM Tris-HCl pH 8.4, 40 mM NaCl. The sample was concentrated and injected onto a sephacryl 16/60 S500 gel filtration column equilibrated in 25 mM Tris-HCl pH 8.4, 50 mM NaCl. A typical elution profile for Vipp1 consisted of three peaks containing 1) Vipp1 superstructures such as filaments eluting at ∼ 40mL within the column size exclusion limit or void volume, 2) Vipp1 rings eluting at ∼ 65 mL, 3) non-polymerised low molecular weight Vipp1 species consistent with monomer or dimers, MBP and TEV eluting at ∼100 mL. Fractions from peak 1 and 2 were pooled and concentrated up to 1 mg/mL. Where necessary, the sample was gel filtrated a second time to reduce residual MBP or TEV contamination. LC-MS/MS confirmed the identity of the Vipp1 band identified by SDS-PAGE. Note that native Vipp1 migrates at ∼38 kDa and not at its expected molecular weight of 28.7 kDa. As Vipp1*Δα*5/6_1-191_ gel filtrates only in peak 3, the removal of MBP and TEV was necessary for a clean SDS-PAGE analysis. An additional affinity chromatography step was therefore included directly before gel filtration using 2 × 5 mL HisTraps (GE Healthcare). As both the MBP and TEV proteins incorporate a hexahistidine tag, the flow through containing Vipp1*Δα*5/6_1-191_ was collected for subsequent steps.

#### Dynamin cloning, protein expression and purification

The rat *dynamin 1* gene (Uniprot code P21575) with the PRD domain truncated was cloned into pOPTM vector yielding an N-terminal MBP fusion with a TEV cleavage site in the linker. A C-terminal hexahistidine tag was included on Dynamin 1. *E. coli* thioredoxin was inserted at the Dynamin 1 N-terminus through Gibson assembly. The role of the thioredoxin, which was not cleaved off, was to stop the large-scale clumping of Dynamin 1 filaments in solution and to facilitate a broad distribution of filaments on the EM grid. Transformed *E. coli* BL21 (DE3) cells were grown on selective LB-agarose plates with ampicillin (100 μg/ml). 2xYT media was inoculated and cells grown at 37 °C until induction at OD600 = 0.6 with 1 mM isopropyl β-D-1-thiogalactopyranoside (IPTG). Cells were grown for ∼15 h at 19 °C and shaken at 220 rpm. All the further steps were carried out at 4 °C unless otherwise specified. For purification, pellets were re-suspended in 50 mM Tris-HCl pH 8.0, 500 mM NaCl, 2 mM DTT, 2 mM EDTA and sonicated on ice. The lysate was clarified by centrifugation in a Ti45 rotor (Beckman Coulter) at 98,000 × g at 4 °C for 45 min and the supernatant loaded onto a self-packed column with ∼10 ml of amylose resin (NEB) pre-equilibrated in 50 mM Tris-HCl pH 9.0, 500 mM NaCl, 1 mM DTT, 1 mM EDTA and 20% glycerol (wash buffer). The resin was washed with 200 mL of wash buffer and the MBP-Dynamin 1 eluted with wash buffer supplemented with 15 mM maltose. The sample was incubated overnight at room temperature with TEV and the products separated by gel filtration using a HiPrep 26/60 Sephacryl S300 column in buffer 20 mM Tris-HCl pH 9.0, 1mM EDTA and 1 mM DTT. Fractions of Dynamin 1 were concentrated to ∼10 mg/ml, flash frozen in liquid nitrogen and stored at -80 °C.

#### Cryo-EM sample preparation and data collection

Under cryogenic conditions *N. punctiforme* Vipp1 exhibits significant preferred orientation so that essential side views for determining the structure were absent. To solve this issue, preformed Dynamin 1 filaments with 37 nm diameter were mixed with Vipp1 prior to vitrification. Rat Dynamin 1 filaments appear equivalent to human Dynamin 1 filaments in the super constricted state (*22*). The Dynamin 1 filaments formed a network on the grid that helped to maintain ice thickness around Vipp1 rings so that positioning at the air-water interface was reduced and side views captured. 37.5 µM Dynamin 1 was incubated with 20 mM HEPES-NaOH pH 7.2, 50 mM NaCl, 1 mM DTT, 2mM GMPPCP and 5 mM MgCl_2_ at room temperature for 2 hours to form filaments. 35 µM Vipp1 was then mixed with the pre-formed dynamin filaments. 4 μl of the mixture was incubated for 30 s on glow discharged holey R2/2 Quantifoil grids before vitrification in liquid ethane using a Vitrobot Mark IV (FEI). Data were collected at 300 kV on a Titan Krios (M02 beamline at eBIC Diamond, UK) equipped with a Gatan Quantum K2 Summit detector. 3206 movies were acquired at a magnification of ×35,971 yielding 1.39 Å/pixel using EPU software. Defocus was between -1.25 and -3.0. Movies were dose-weighted over 40 frames with 10 s exposures. Total dose was 50 e/Å^2^.

#### Cryo-EM image processing

Individual movie frames were aligned with MotionCor2 (*55*) and the contrast transfer function estimated using CTFFIND4 (*56*). All subsequent processing was carried out using Relion 3.0 (*57*). Particles were picked manually to generate initial 2D class averages that were subsequently used for reference based auto-picking. Extracted particles were subjected to five rounds of 2D classification resulting in a cleaned stack of 109,715 particles. To generate the initial 3D model, a subset of 2D classes comprising C14 symmetry top views and side views with diameter range from 28 nm to 32 nm were selected (24,355 particles). 3D classification was carried out in C1 using a featureless hollowed cylinder as the initial reference. One class containing 9,991 particles yielded a ring with distinct C14 symmetry, which was chosen for high-resolution reconstruction first. These particles were then 3D autorefined with C14 symmetry applied reaching 8.5 Å resolution. The resulting map (Intermediate map 1) showed clear secondary structure features and was used as the new C14 reference volume for a second round of processing. In round two, side views only of 2D class averages were selected and 3D classified iteratively in C1 using Intermediate map 1 as a reference volume. In this way 15,767 side views with C14 symmetry were isolated. These side views were combined with 3,663 C14 top view particles obtained during 2D classification. A 3D autorefinement was undertaken with C14 symmetry applied to yield a reconstruction at 7.0 Å resolution (Intermediate map 2). Individual particles were then corrected for beam-induced motion for a third round of processing. One round of 3D classification was undertaken in C1 using Intermediate map 2 as the reference volume. A final stack of 17,114 particles was then used for 3D autorefinement with C14 symmetry applied reaching 6.8 Å resolution. Post-processing yielded 6.5 Å resolution with an auto-estimated B-factor (*58*) of -291.9 Å^2^ applied to sharpen the final 3D map. Resolutions reported are based on gold standard Fourier shell correlations (FSC) = 0.143. Once the Vipp1_C14_ structure was built and targeted masks of asymmetric units or individual rungs could readily be generated, multiple subtraction based local refinements including symmetry expansion strategies were attempted but no improvement in resolution was observed. A similar strategy as implemented for Vipp1_C14_ was carried out to generate all other ring symmetries including Vipp1_C11-C13_ and Vipp1_C15-C17_. A B-factor of -320 Å^2^ was applied to these maps. The hand of the electron density maps was unambiguously determined by fitting the PspA crystal structure (PDB code 4WHE), which has a distinct axial twist and asymmetry. Statistics for data collection and 3D refinement for all maps are included in Table S1.

#### Model building and refinement

Rung 3 of the Vipp1_C14_ structure was built first. A secondary structure prediction was obtained using Psipred (*46*). A partial Vipp1 homology model based on PspA (PDB code 4WHE) was generated using I-Tasser (*59*). The model was trimmed to include amino acids 24-142, which represents the hairpin motif. Importantly for obtaining an accurate sequence register in the Vipp1 structure, Vipp1 aligns robustly with PspA in this region with 32.5 % sequence identity, 59 % similarity and crucially 0 % gaps. The hairpin readily fitted into the Vipp1_C14_ map requiring only minor adjustments. Overall the hairpin from Vipp1_C14_ rung number 3 (PDB 6ZW4) and PspA hairpin motifs have a C*α* RMSD = 2.2 Å (Fig. S4E). The hairpin homology model provided an important anchor for subsequently building the N-terminal helix *α*0 and C-terminal helices *α*4 and *α*5. The resolution for the bulk of Vipp1_C14_ within rungs 3 and 4 was ∼ 5 Å so that the main chain could be easily traced and helices *α*0-*α*5 clearly assigned and built using COOT (*60*). Significant attention was paid to regions of high sequence conservation as a guide for sequence register within predicted interfaces. Similarly, our co-evolutionary contact maps (Fig. 1B) were used to confirm interfaces and expected sequence register. Ultimately, the accuracy of the sequence register was experimentally assessed by the introduction of a cysteine pair within Interface 3 and tracking cross-links (Fig. S6A). Rosetta (*61*) was used to improve the geometry. The subunit from rung 3 was copied and rigid body fitted into all other rungs within the asymmetric unit using Chimera Fit in map command (*62*). COOT and ISOLDE (*63*) were used to adjust for rung specific conformational changes. Using PHENIX (*64*), non-crystallographic symmetry (NCS) was applied to each asymmetric unit to generate a complete Vipp1_C14_ 84 chain model. This model was truncated to main chain and rigid body and B factor refined in PHENIX. For all other ring symmetries, the Vipp1_C14_ asymmetric model was fitted using Chimera Fit in map command. COOT and ISOLDE (*63*) were used to adjust for conformational changes specific to ring symmetry. For each ring, NCS was applied to generate complete ring models. All subsequent steps were as for Vipp1_C14_. The final models were assessed using Molprobity and statistics outlined in Table S1 (*65*). The correlation between map and model (CC_mask_) as generated by the phenix.map_model_cc command was C11- 0.8, C12- 0.82, C13- 0.77, C14- 0.84, C15- 0.8, C16- 0.67, C17- 0.58.

#### Elastic network model

To understand the relationship between inter-rung stacking and the creation of domed curvature in Vipp1 rings, molecular modeling was used focused on the smallest ring system-Vipp1_C11_ for simplicity. Each residue was represented by a single bead at the position of the C*α* atom. The potential energy of the structure was defined by an elastic network model (ENM), meaning that interactions between residues nearby in the experimental structure were restrained by harmonic springs. Nearby in the contact map was defined by C*α* distances within 10 Å. All springs were given the same stiffness.

The elastic network for each rung was identical, both internally (intra-rung) and in the interactions made with the rungs above it and below it (inter-rung). While the experimental Vipp1_C11_ structure enforced each monomer to be identical within a rung, the monomers between rungs showed small differences. To create the intra-rung network, the contact map for all monomers in rungs with a complete structure (11*4 monomers in Vipp1_C11_ rungs 2-5) were compared and a spring was created for each contact provided it existed in >50 % of the monomers. As the conformational changes in the monomers between different rungs are small, only a few contacts were removed in this process. Those removed were localized to the regions showing the largest shifts between rungs, namely Hinges 2 and 3 within the Vipp1 monomer (Fig. 4B). Removing these outlier contacts allows the network to better model the inherent flexibility of the Vipp1 monomer. Note that the intra-rung contacts included contacts between monomers within the same rung. The natural length of each spring was defined as the average of its contact distances over the monomers. In this way, an ENM for an average rung was created. The average rung best matched rung 3, with a C*α* RMSD of 0.5 Å. The inter-ring contact map was defined by the interactions between Vipp1_C11_ rungs three and four (Fig. 2B) as rung 5 at the bottom is incomplete.

We then computed the equilibrium structure for different stack sizes by minimizing the elastic energy. This minimization was performed by molecular dynamics (MD) at a low temperature followed by a steepest decent minimization. SMOG2 (*66*) with the template “ENM” was used to create topology files for the MD software GROMACS (*67*) using the Vipp1_C11_ PDB structure as input (PDB code 6ZVR). These topology files were processed as described in the previous paragraph. Initial structures for minimization were created in VMD by manually copying rung three and translating it N times, where N is the desired number of rungs.

The equilibrium structures resulted from balancing the competing effects of 1) the inter-rung interactions driving curvature and 2) the geometrical constraints of doming. While in principle some of the strain could be alleviated by breaking links, this is not allowed in the ENM. This constraint is not present in the experimental system, which may explain why the densities show that parts of the upper and lower rungs do fail to assemble properly and become disordered. The rotations between rungs in the equilibrium structures were analyzed by measuring the angle formed by helix *α*5 with the dome axis (Fig. 5B). The axis is defined by the z-axis in the experimental structure. The line along the direction of helix *α*5 is defined by two points taken as the centers of mass of residues 194-202 and residues 211-219.

#### Vipp1 cysteines crosslinking assay

DTT was removed from 1 mg/mL Vipp1_L86C_, Vipp1_L193C_ or Vipp1_L86C/L193C_ using PD MiniTrap G-25 columns (GE Healthcare) equilibrated in 25 mM Tris-HCl pH 8.4, 50 mM NaCl. Samples were diluted to 5 µM and incubated with either 10 mM DTT, ortho-Cu(II)1,10-phenanthroline (CuP, stock 10 mM in 20 % ethanol) or 5 µM 1,4-Butanediylbismethanethiosulfonate (MTS-4, stock 50 mM in chloroform) for 1 h at room temperature. Vipp1_L86C/L193C_ cross-linked samples were rescued with 10 mM DTT for 1 h at room temperature. Non-reacted cysteines were blocked by the addition of 10 mM N-ethylmaleimide (NEM, stock 0.5 M in 100 % ethanol). Samples were evaluated by SDS-PAGE.

#### Liposome preparation

Liposomes were prepared using *E. coli* total lipid extract (Avanti polar lipids, US). Lipid extract was dissolved in chloroform at 25 mg/mL in a glass vial (Thermo Fisher Scientific). Chloroform was evaporated and the lipid dried for 1 h in a vacuum desiccator. The residual lipid film coating the bottom of the vial was hydrated using liposome reaction buffer (20 mM HEPES, pH 8.0 and 80 mM KCl) at a concentration of 6 mg/ml. The lipid was resuspended by vortexing and gentle sonication with a needle tip for 2 min on ice. The suspension was extruded through polycarbonate membranes with 1 or 0.2 µm pore size using a mini-extruder (Avanti Polar Lipids) to create large or small unilamellar vesicles (LUV/SUV). LUV and SUV were stored at 4 °C for subsequent use.

#### Vipp1 liposome binding assays and negative stain EM

Liposome binding assays were performed by incubating freshly prepared 2 mM SUV with and without purified 5 µM Vipp1 for 2 h at room temperature in liposome reaction buffer. 5 μL of each sample was loaded onto glow-discharged 200-mesh carbon coated copper grids and stained with 2 % uranyl acetate (UA). Images were acquired using a FEI Tecnai Spirit microscope equipped with a 2 K Eagle camera.

#### Spin Assay

To detect Vipp1 membrane binding a spin assay was used. 10.5 µM Vipp1 was ultra-centrifuged at 50,000 x g at 20 °C for 15 min using a TLA100 rotor to remove any initial aggregation. The supernatant from this first spin was collected and incubated with and without 2 mg/ml LUV for 1 h at room temperature. Samples were subjected to a second spin at 30,000 x g at 20 °C for 30 min. The pellet (P) and the supernatant (S) were harvested, made up to equal volumes in LDS sample buffer and analysed by SDS-PAGE.

#### Vipp1 monolayer assays

Lipid monolayers were prepared using *E. coli* total lipid extract (Avanti Polar Lipids). A custom-made Teflon block containing 4 mm x 4 mm diameter wells were filled with 50 μL of assay buffer (20 mM Tris-HCl, pH 8 and NaCl 50 mM). A 5 μL drop of 0.1 mg/ml lipid dissolved in chloroform was applied to the top of the buffer solution and the chloroform left to evaporate for 1 h. A non-glow discharged carbon coated copper grid was gently placed on top of the lipid layer with the carbon side faced towards the lipid layer. Subsequently, 14 µM Vipp1 was injected into the well using a side port. The control wells containing either monolayer (no protein) or protein only (a drop of chloroform but no lipid) were set up in parallel. Samples were incubated for 2 h before grids were recovered and immediately stained with 2 % UA and imaged using a FEI Tecnai Spirit microscope equipped with a 2 K Eagle camera. Images were taken at 89020 x magnification and 3.37 Å pixel size. 263 and 156 images were collected of Vipp1 with and without lipid monolayer, respectively, as described above. Gctf1.06 (*68*) was used for estimating the contrast transfer function. Processing steps including particle picking and extraction, and 2D classification were carried out using Relion 3.1 (*57*). The final class averages were generated from stacks comprising 11,972 and 2,165 particles for Vipp1 with and without lipid monolayer.

